# Ploidy and neuron size impact nervous system development and function in *Xenopus*

**DOI:** 10.1101/2025.04.22.650131

**Authors:** Xiao Liu, Christine Wan, Sara Aijaz Shah, Rebecca Heald

## Abstract

Neuron size varies significantly over evolution, contributing to diverse nervous systems of variable complexity, while aberrant neuron size is associated with neurodevelopmental and degenerative diseases. How do neuron cell body and neurite size and organization impact nervous system development and function? To systematically study effects of neuron size on the vertebrate nervous system, we characterized triploid *Xenopus* tadpoles that possess a 1.5-fold increase in genome size compared to diploids. Triploids displayed a linear increase in total neuronal volume and a superlinear increase in membrane surface area. Imaging, flow cytometry, and RNA-seq analyses revealed that triploid brains were morphologically and transcriptionally similar to diploid brains, but less proliferative, containing fewer neurons and displaying increased global activity. Interestingly, physiological differences at the neuron and nervous system levels affected swimming behavior in tadpoles. Our findings thus establish a framework to link genome size, neuron size, and nervous system development and function in vertebrates.

## INTRODUCTION

Cells vary dramatically in size; yet the size of a particular cell type is relatively constant and mechanisms are in place to ensure that cell size is properly tuned^1,2^. Previous studies indicate that cell size impacts many aspects of cell biology, including rates of biosynthesis, metabolic activity, and response to stimuli^2^. Nevertheless, the physiological consequences of cell size remain poorly understood, particularly in the brain.

Of the two main building blocks of the nervous system, neurons and glia, neurons display much larger variations in cell size over evolution^3–5^. Although underlying mechanisms are not entirely clear, one cell size scaling factor is genome size. In amphibians, where interspecies hybridization and spontaneous polyploidization have resulted in dramatic genome size variation, a positive correlation between neuron cell body size and genome size has been demonstrated^6,7,8^. In *Drosophila*, fish, and mammals, somatic polypoid neurons are also associated with a larger cell body^9–11^. However, previous studies have not elucidated the effects of genome size on neuron size beyond cell body size. While the neuronal cell body (soma) supports cellular housekeeping functions and affects overall membrane potential kinetics, the neurite compartment (axon and dendrites), which contributes significantly to total cell volume, critically impacts signal propagation and connectivity. Therefore, size and shape parameters of both the cell body and neurite compartments, including cell body size, neurite length and diameter, and neurite network complexity (Figure 1A), should be examined collectively to understand the impact of neuron size on nervous system function.

**Figure 1.**
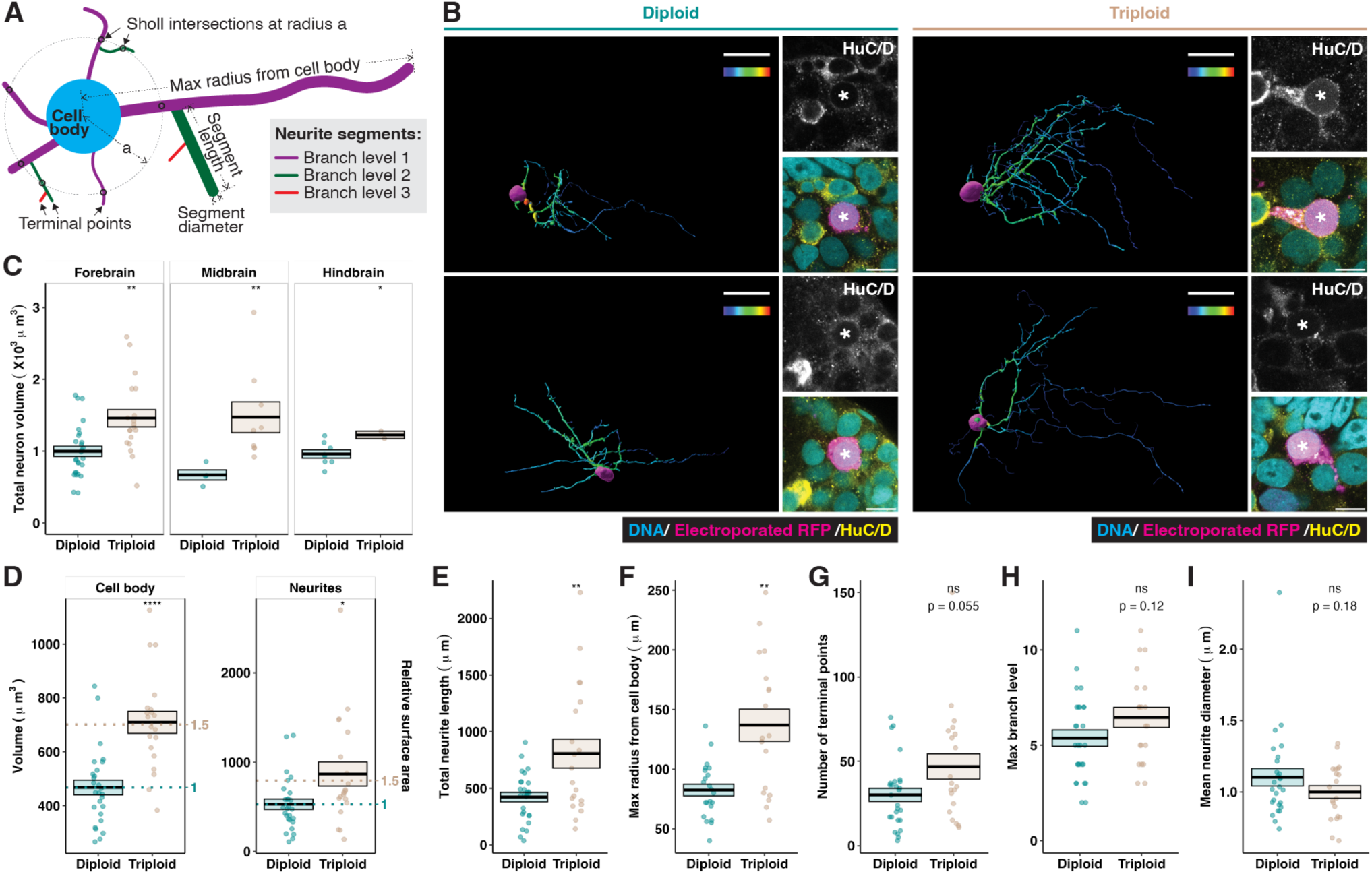
Triploid neurons are larger in volume and extend longer neurites. **A.** Schematic showing different neuron size and shape parameters. **B.** Two representative examples of reconstructed electroporation-labeled diploid and triploid neurons with right panels showing higher magnification micrographs of their cell bodies. The reconstructions were rotated to show the longest axes of the neurons, and therefore cell bodies are not in the same orientation. Immunostaining of a pan-neuronal marker, the RNA-binding Hu proteins HuC/D (yellow), was used to confirm the neuronal identity of the labeled neurons^82^ (magenta). Micrographs were taken at different laser settings, their color channels adjusted separately (see STAR methods). In the reconstructions, scale bar, 30 μm; color bar, segment mean diameter 0-3 μm. In the zoomed-in micrographs, scale bar, 10 μm; asterisks, cell body of the labeled, reconstructed neuron. **C.** Comparison of total neuron volume between diploid and triploid neurons. Numbers of diploid/triploid neurons measured were 26/17 in the forebrain, 4/8 in the midbrain, and 8/2 in the hindbrain. See also Figure S2A for sholl analysis of these neurons. Only forebrain neurons were used for analyses in D-I. **D.** Comparison of cell body and neurite volume. Dotted lines, 1- and 1.5-fold of diploid mean. **E-I.** Comparison of various size parameters: total neurite length (E), maximum radius from the cell body (F), number of terminal points (G), maximum branch level (H, see also Figure S2B), and mean neurite diameter (I, see also Figure S2C). In C-I, each dot represents one neuron. Crossbars denote mean ± SEM. *, p<0.05; **, p<0.01; ****, p<0.0001; ns, not significant, t test.

Variation in any neuron size parameter could alter brain activity. Increased neurite diameter correlates with faster action potential kinetics^12^, while increased neurite length could mean a more extensive or expansive neurite network, leading to greater information capacity and connectivity potential^12,13^. Additionally, larger neurons with increased membrane surface area have altered capacitance and resistance, and potentially harbor more ion channels and synaptic connections, resulting in reduced noise and higher temporal precision in electrical activity^12,14–17^. These properties may confer certain advantages to larger neurons. Indeed, the expansion in brain size across mammalian evolution is accompanied by a significant increase in average neuron mass^18^. In humans, larger pyramidal neurons are associated with more efficient information relay and higher IQ scores^14^. Conversely, a downward shift in neuron size is observed in the brains of patients with neurodegenerative diseases including Alzheimer’s Disease^19^, Huntington’s Disease, and Schizophrenia^20^. However, evolution does not simply select for larger neurons. In amphibians, brain complexity correlates inversely with neuron cell body size^7,8^. In birds, relative brain size decreases as genome size and cell size increase^21,22^. Additionally, increased neuron size is implicated in neurological disorders such as Lhermitte-Duclos disease and tuberous sclerosis^23–25^. Thus, our understanding of the effects of neuron size on the brain is primarily anecdotal and more systematic approaches are needed.

In this study, we leverage *Xenopus*, a powerful model to study development and function of the vertebrate nervous system^26^ as well as biological size control mechanisms^6,27,28^, to examine the links between genome size, neuron size, and neurodevelopment. By comparing individual neurons in diploid and triploid *Xenopus* embryos, we reveal how different parameters of neuron size scale with genome size. Combining imaging and flow cytometry approaches, we construct a framework to characterize how neuron size changes impact the development and function of the *Xenopus* brain. We show that triploid brains are wired with fewer, larger neurons and display a global change in neuronal activity, ultimately linking physiological changes at the neuron and brain levels to distinct functional behaviors at the organism level.

## RESULTS

### Triploid neurons are larger than diploid neurons

To generate *X. laevis* triploid embryos, we adapted established procedures^29^ and cold shocked half of each clutch of eggs shortly after fertilization to block extrusion of the second polar body, resulting in the inclusion of two sets of maternal chromosomes in addition to one set of paternal chromosomes (Figure S1A). Consistent with previous studies^27^, triploid embryos developed at a similar rate as their diploid siblings, did not display physical abnormalities, and developed into normal-looking adult frogs (Figure S1A-C). Flow cytometry analysis of dissected, dissociated brains stained with Hoechst DNA dye confirmed the ploidy increase in triploid brains. Triploid DNA content showed a marked rightward shift to around 1.5-fold of that of diploids (Figure S1D). Consistent with this, *in toto* imaging revealed a 1.5-fold increase in nuclear volume in triploid neurons compared to diploids (Figure S1E).

Across numerous cell types in all three domains of life, cell size scales positively with genome size^30,31^. Likewise, epithelial cells of experimentally induced triploid *Xenopus* tadpoles were observed to be approximately 50% larger than those of diploids^27^. However, given their highly complex morphology, neuron size is much more difficult to assess. To determine how different parameters of neuron size (Figure 1A) scale to a change in genome size, we performed sparse neuron labeling to profile the size of single neurons *in vivo*. We microinjected fluorescent protein-encoding plasmids into brain ventricles of developing *X. laevis* embryos and used targeted electroporation to facilitate uptake by periventricular neuron progenitors^32^, which gave rise to labeled neurons that migrated into the developing brain layers^33^. Brains with single-labeled neurons were then cleared and imaged *in toto* with refractive index matching, and the imaged neurons were reconstructed in 3D (Figure 1B).

Pooling neurons by brain region, we found that total neuron volume (including cell body and neurites) was significantly greater in triploids compared to diploids (Figure 1C). We were best able to obtain images for whole, individually labeled neurons in the forebrain, where Sholl analysis comparing neurite complexity revealed the highest consistency between diploid and triploid neurons (Figure S2A), indicating that the labeled neurons belonged to similar neuronal subtypes by morphological measures. We therefore used forebrain neurons for detailed size profiling.

The increase in total volume of triploid neurons resulted from changes in the cell body and neurite compartments, with both scaling approximately linearly with the 1.5-fold ploidy change (Figure 1D). Triploid neurites were significantly longer, with the total neurite length per neuron nearly double that of diploids (Figure 1E). The increased length was distributed to expand the radius of the neuron (Figure 1F), rather than to fill in the radius with more extensive arbors. Neither the number of terminal points (Figure 1G) nor branch levels (Figure 1H) changed significantly in triploid neurons, with neurite length similarly distributed over branch levels in both ploidies (Figure S2B). These results indicate that triploid neurons do not possess distinct branching complexity or patterns compared to diploids, despite increased neurite length and volume. Unlike other size parameters measured, triploid neurite diameter showed a subtle downward shift (Figures 1I and S2C). Thus, triploid neurons scaled up their volume by having a larger cell body and by growing neurites that were longer, but not thicker.

### Neuronal surface area scales superlinearly with ploidy and volume

When averaged, the decrease in neurite diameter in triploid neurons was not statistically significant (Figure 1I). However, closer examination of how neurite length was distributed over different diameters revealed that a larger proportion of triploid neurite length possessed smaller diameters (Figure 2A). This resulted in an increase in total neuronal cell surface area in triploids compared to diploids. Compared to the neuronal cell body that can be roughly modeled as a sphere in which surface area increases sublinearly with volume (Figure 2B, top panel), neurites resemble thin tubes, for which surface area increases superlinearly with volume if the diameter decreases as the volume increases (Figure 2B, bottom panel). Consistent with these simple geometric predictions, we observed that while the volume of cell body and neurite compartments scaled linearly with the 1.5- fold ploidy change (Figure 1C), the surface area of the cell body scaled sublinearly, and the surface area of the neurites scaled superlinearly (Figure 2C, left and middle panels). Therefore, although cell volume was divided between the cell body and neurite compartments in a similar ratio in both ploidy neurons (Figure 2D, left panel), cell surface area was not. A higher percentage of cell surface area was contributed by neurites in triploid neurons (Figure 2D, middle panel), where the differential scaling of surface area to volume ratio between neurites and cell body was significantly more dramatic (Figure 2D, right panel). Since neurites contributed ∼5X more surface area than the cell body (Figures 2C-D and S3), total surface area followed the trend of neurite surface area and scaled superlinearly with ploidy and cell volume (Figure 2C, right panel).

**Figure 2.**
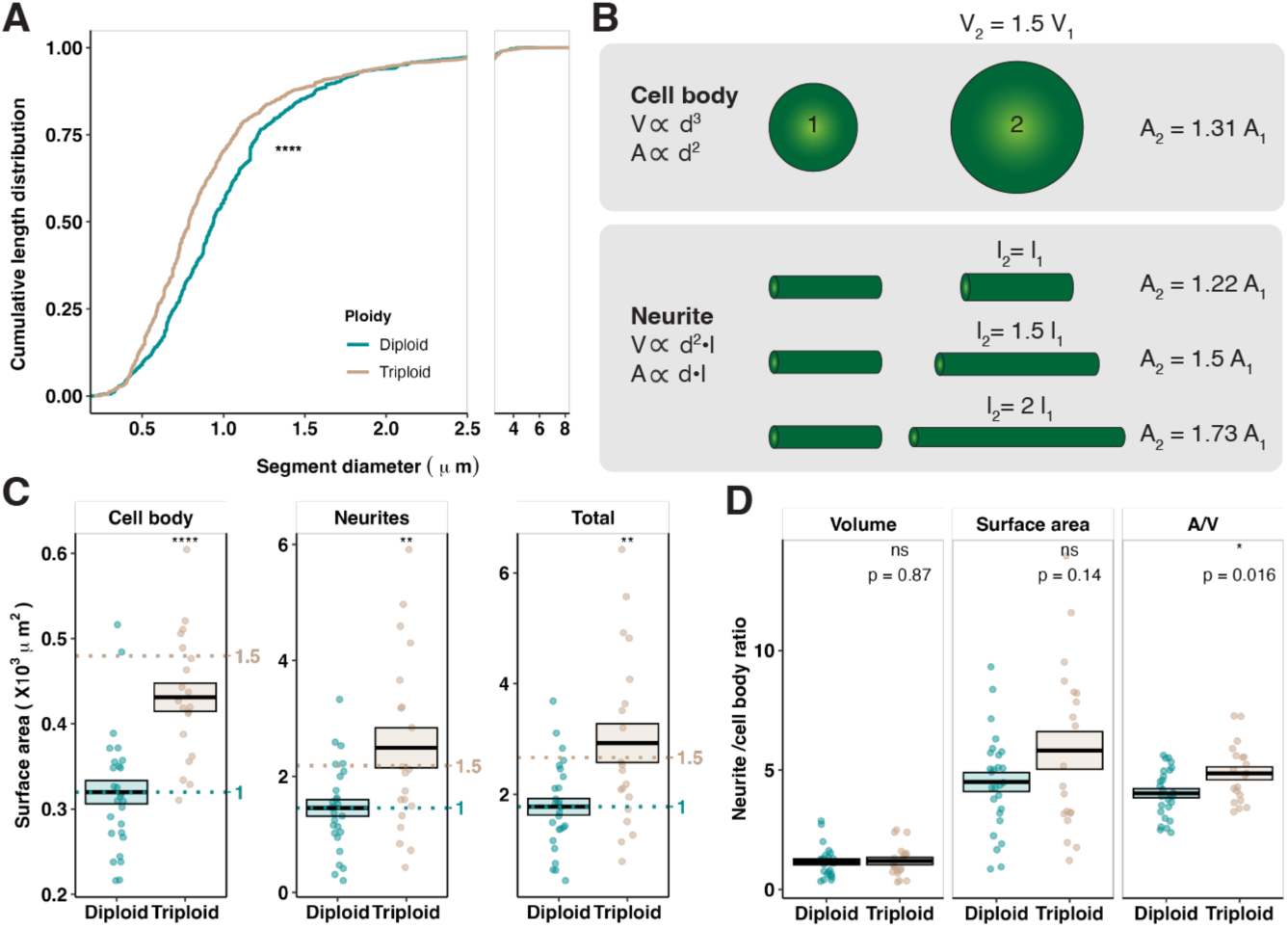
Cell surface area scales superlinearly with neuron ploidy and volume. **A.** Cumulative length distribution of diploid and triploid neurites at different diameters. Leftward shift indicates a larger portion of neurite length with smaller diameter. ****, p<0.0001, Kolmogorov– Smirnov test. **B.** Diagram modeling differential increase in volume (V) and surface area (A) in relation to diameter (d) and length (l) in the cell body and neurites. **C.** Comparison of cell body, neurite, and total surface area between diploid and triploid neurons. Dotted lines, 1- and 1.5-fold of diploid mean. **D.** Ratios between the neurite and cell body compartments for volume, surface area, and surface area/volume (A/V) in diploid and triploid neurons. See also Figure S3. In C-D, each dot represents one brain. The same 26 diploid and 17 triploid forebrain neurons as in Figure 2 were analyzed. Crossbars denote mean ± SEM. *, p<0.05; **, p<0.01; ****, p<0.0001; ns, not significant, t test.

### Triploid *Xenopus* tadpoles possess normal brain morphology

Interestingly, despite the significant increase in ploidy and neuron size, triploid brains displayed largely unaltered morphology (Figure 3A), with a similar aspect ratio (Figure 3B) and proportions of forebrain, midbrain, and hindbrain compared to diploid brains (Figures 3C and S4A-C). Brain size increased in triploids but only with a small effect size disproportionate to the ploidy increase (Figure 3D). This change in brain size was independent of the sex of the tadpole (Figure S4D).

**Figure 3.**
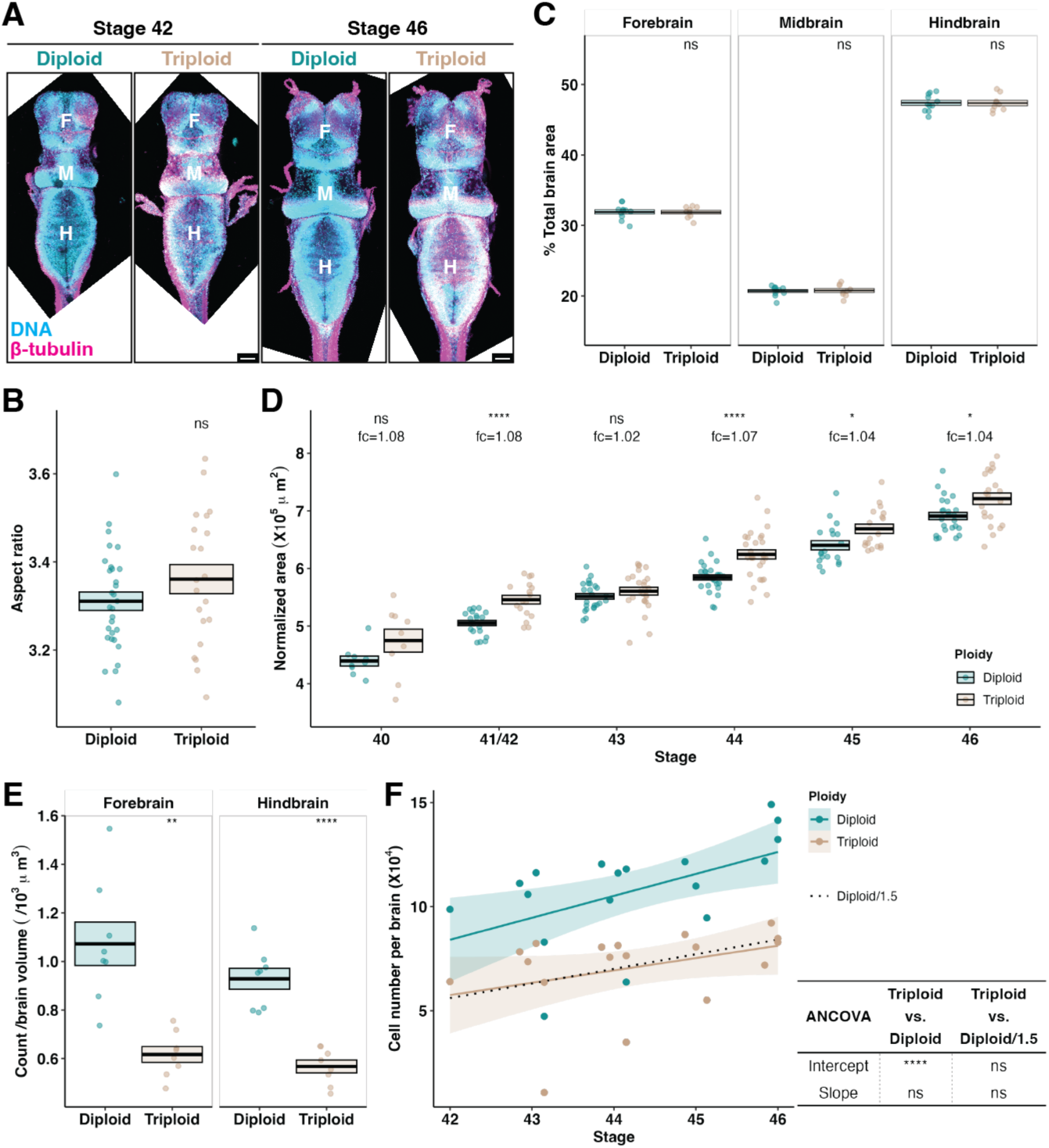
Triploid brains are morphologically similar to diploid brains, but contain significantly fewer cells. **A.** Representative z-projected micrographs of diploid and triploid brains at the indicated developmental stages. Stage 46 images were stitched from two overlapping tiles. F, forebrain; M, midbrain; H, hindbrain. Scale bar, 100 μm. **B.** Aspect ratio of diploid and triploid brains at stage 46. Each dot represents one brain and a total of 30 diploid and 21 triploid brains from 4 independent clutches were examined. **C.** Proportion of the indicated brain region in diploid and triploid brains at stage 46. Each dot represents one brain and a total of 18 diploid and 15 triploid brains from 3 independent clutches were examined. See also Figures S4A-C for data from different developmental stages. **D.** Size comparison of diploid and triploid brains across multiple developmental stages. Each dot represents one brain. Numbers of diploid/triploid brains examined were 9/9 at stage 40, 20/17 at stage 41/42, 24/26 at stage 43, 30/28 at stage 44, 19/18 at stage 45, and 24/21 at stage 46. Brains were from 4 independent clutches. Areas were normalized to adjust for clutch variance. See also Figure S4D for how sex did not impact brain size. In B-D, crossbars denote mean ± SEM. *, p<0.05; **, p<0.01; ***, p<0.001; ****, p<0.0001; ns, not significant, t test. See also Figures S4E-G for similar metrics in *X. tropicalis*.

Similar observations were made in *X. tropicalis*, another *Xenopus* species with a smaller genome size. In artificially induced *X. tropicalis* triploids, brain morphology and organization were largely preserved compared to diploids (Figures S4E-F), and the brain size increase, although significant, was small in effect size (Figure S4G).

With larger neurons as building blocks, why are triploid brains not much larger? With imaging-based cell counting (Figure 3E) and flow cytometry of dissociated brains (Figure 3F), we observed a striking decrease in cell number in triploid brains. This 1.5-fold decrease was significant and consistent across multiple brain regions and developmental stages, thereby offsetting the neuron size increase and resulting in only a minor increase in triploid brain size.

### Triploid brain cells are less proliferative but do not exhibit differences in transcription

To investigate why triploid brains contained significantly fewer cells, we performed DNA content-based cell cycle analysis on cells dissociated from tadpole brains. We found that triploid brains contained a significantly smaller proportion of G_2_/M cells than diploids (Figure 4A), indicating reduced proliferation. We then analyzed two cell cycle markers, proliferating cell nuclear antigen (PCNA), an indicator of DNA replication^34,35^, and phospho-histone 3 (pH3), which labels condensing DNA during M phase^36,37^, by immunofluorescence (Figure 4B) and flow cytometry. We observed distinct PCNA expression patterns in tadpole brains, which we classified as PCNA negative (G_0_), PCNA diffuse, and PCNA punctate (S), respectively (Figures 4C). pH3 staining was positive only in mitotic cells that contained condensed chromosomes (Figure 4D).

**Figure 4.**
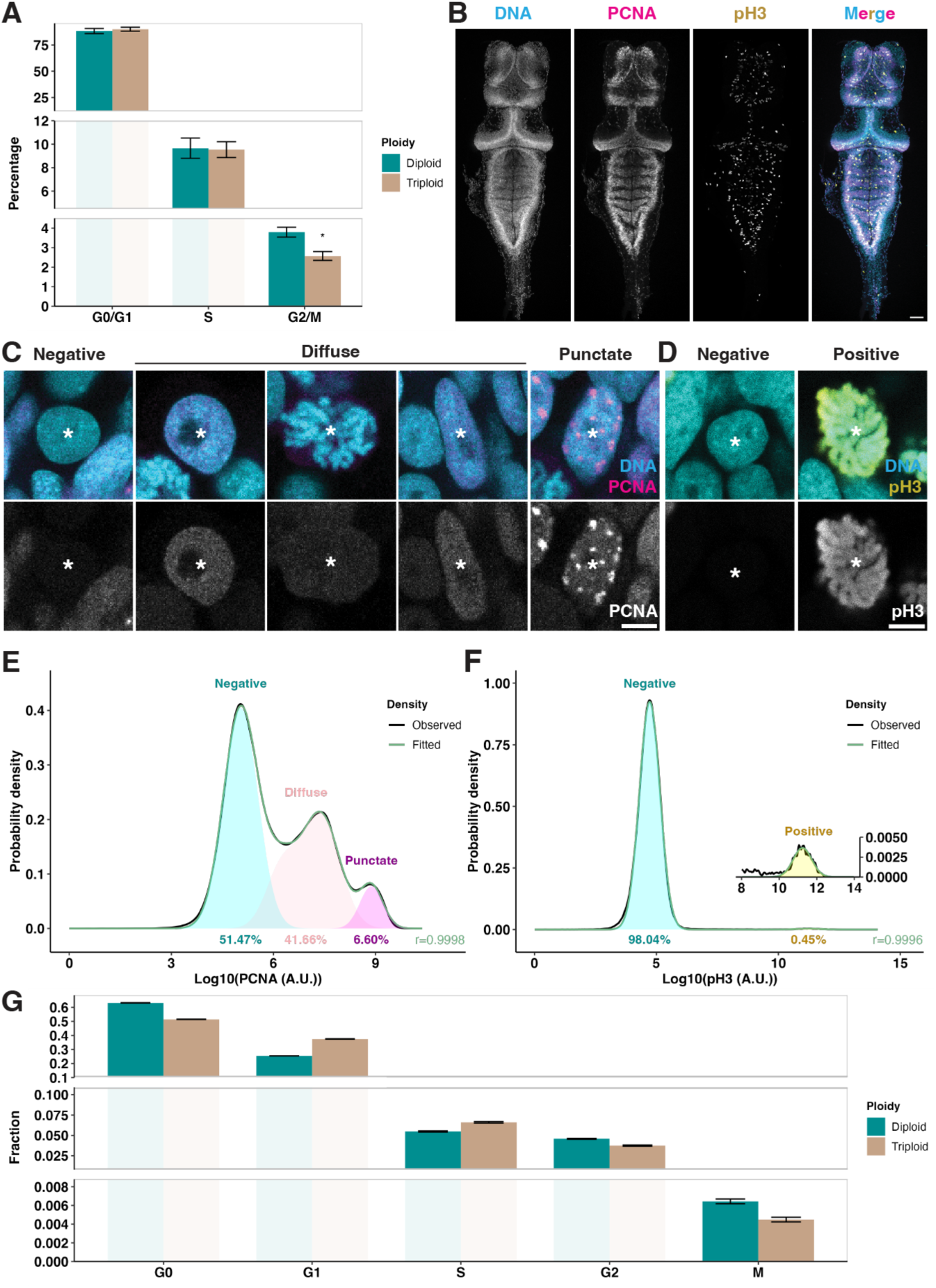
Triploid brains are less proliferative than diploid brains. **A.** Cell cycle analysis based on Hoechst intensity using the Dean Jett Fox (DJF) model^83^. Data were pooled from flow cytometry runs of 5 pairs of clutch-controlled diploid and triploid samples, each containing 5-6 developmental stage 46 brains. Mean ± SEM are shown. *, p<0.05, paired t test. **B.** Representative z-projected micrograph of a dissected stage 46 brain after co-staining with Hoechst and antibodies against PCNA and pH3, before dissociation for flow cytometry. Image was stitched from two overlapping tiles. Scale bar, 100 μm. **C.** Representative micrographs of three different PCNA expression patterns in the brain: 1) PCNA negative (G_0_) nuclei were all round-shaped, neuronal nuclei; 2) A diffuse PCNA pattern was observed in various nucleus shapes and phases, including round neuronal nuclei, elongated neural progenitor nuclei, and mitotic nuclei containing condensed chromosomes; 3) PCNA punctate (S) nuclei all possessed an elongated shape typical of progenitors. Asterisks, nuclei. Scale bar, 5 μm. **D.** Representative micrographs of two different pH3 expression patterns in the brain. Asterisks, nuclei of interest. Scale bar, 5 μm. **E.** Probability density distribution of Log10 PCNA level of a representative flow cytometry sample of dissociated brain cells at developmental stage 46. Gaussian distributions were used to fit the three populations as shown in C. See also Figure S5A-C for DNA content distributions of the three PCNA populations. **F.** Probability density distribution of Log10 pH3 level of a representative flow cytometry sample of dissociated brain cells at developmental stage 46. Gaussian distributions were used to fit the two populations as shown in E. See also Figure S5D-E for DNA content distributions of the two pH3 populations. **G.** Comparison of fractions of brain cell populations in different cell cycle phases at developmental stage 46. Fractions were calculated from 6 diploid and 5 triploid brains combining DNA content-based cell cycle analysis (as in A) and PCNA/pH3 staining (as in E-F). Error bars mark 95% confidence interval. See also Figure S5F for comparison of cell counts. See also Figure S5G for cell death data and Figure S6 for brain-specific RNA-seq data.

Flow cytometry reliably detected these staining patterns; and combining PCNA and pH3 staining with Hoechst-based DNA content analysis enabled refinement of cell cycle analysis to classify brain cells into each individual cell cycle phase (Figures 4E-F and S5A-E). Compared to diploids, triploid brains contained a significantly larger cell population in G_1_ and S phases, but a significantly smaller population in G2 and M phases (Figure 4G). Triploid brains, which had not only a lower mitotic index (Figure 4G), but also a smaller total cell count (Figure 3F), contained less than half the number of mitotic cells as diploid brains (Figure S5F). In contrast, triploid brains did not show a significant difference in cell death ratio compared to diploid brains by caspase staining (Figure S5G). The decreased proliferation and unaltered cell death in triploid brains indicated that they were less efficient in producing a pool of post-mitotic neurons and glia to make up the brain. Indeed, both the percentage and the absolute number of accumulated G_0_ cells were significantly lower in triploid brains (Figures 4G and S5F). Together, these data account for the decreased cell number in triploid brains and reveal a significant effect of increased ploidy and cell size on cell cycle progression and cell birth/death dynamics.

Despite the altered cell cycle dynamics, triploid brains showed a remarkably similar transcriptional distribution compared to diploid brains (Figures S6), indicating that triploidy, at least within one generation, does not lead to a significant imbalance in gene expression, and that physiological changes observed in triploid brains were at least in part due to post-transcriptional changes caused by increased cell size.

### Triploid brains display a global increase in neural activity

That triploid brains are built with significantly fewer neurons compared to diploid brains indicated wiring and activity differences. With an increased surface area to volume ratio, triploid neurons could possess an altered number and distribution of ion channels, changing their membrane potential control. Additionally, the increased neurite length might lead to altered connectivity.

To determine whether neural activity is impacted in triploids, we imaged brains of freely swimming stage 46 tadpoles and assessed global brain activity by the level of phospho-extracellular signal regulated kinase (pERK) in relation to the level of total ERK. pERK is an established reporter of neural activity downstream of calcium signaling^38,39^. Altered pERK landscape has been reported in animal models of neurological disorders like schizophrenia^39,40^. In tadpole brains most pERK-expressing cells were positive for neuronal markers and had round nuclei typical of neurons rather than elongated nuclei characteristic of proliferating neuron progenitors (Figure S7A-C). Stimulation of tadpoles by tapping their dishes for 15 minutes was sufficient to trigger a significant alteration in pERK/ERK intensity (Figure S7D-E). These observations indicate that pERK/ERK staining reflects neuronal activity rather than changes in cell growth and proliferation.

Compared to diploids, triploid brains showed a significant increase in pERK/ERK levels in all three brain regions (Figures 5A-B and S8A), whereas ERK levels were similar (Figures 5C-D), consistent with our RNA-seq data showing no differential transcription between diploid and triploid brains (normalized by distribution) of either *X. laevis* ERK homeolog (Figure S8B). Together, these data suggest a global elevation in neural activity in triploid brains and support our model that physiological changes in triploid brains occur at the post-transcriptional level.

**Figure 5.**
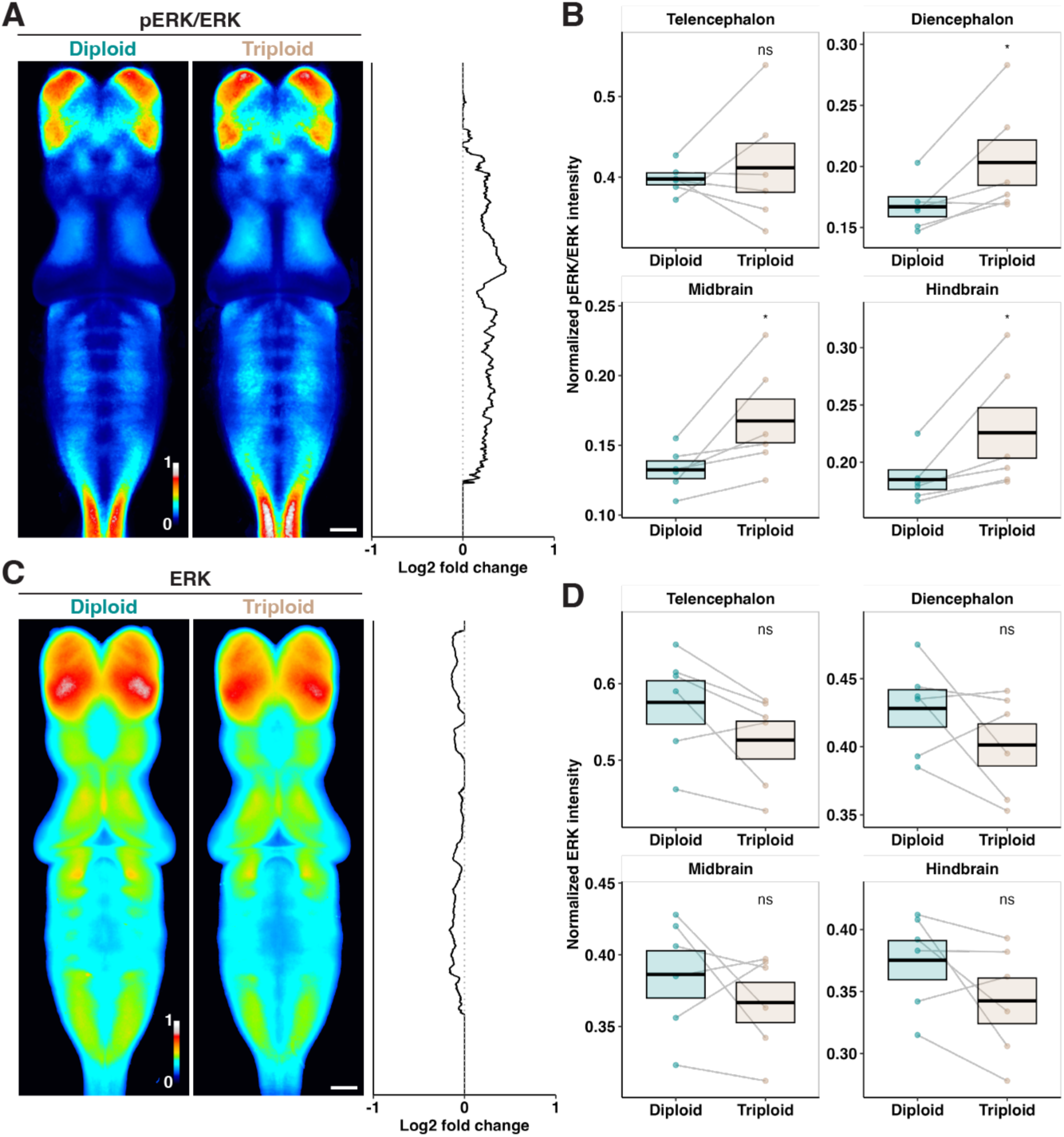
Triploid brains display changes in neural activity. **A.** Heatmap showing normalized, averaged pERK/ERK intensity of 27 diploid and 26 triploid brains of unstimulated, freely swimming stage 46 tadpoles from 6 independent clutches. Log2 fold changes (triploid/diploid) of pERK/ERK intensity along Y axis are plotted to the right. Scale bar, 100 μm. See also Figure S7 for pERK expression patterns and response to stimuli. **B.** Comparison of pERK/ERK intensity in the indicated brain regions. Telencephalon and diencephalon make up the forebrain. Each dot represents the mean value of one clutch. Grey lines connect dots from the same clutch. Crossbars denote mean ±SEM. *, p<0.05; ns, not significant, paired t test. See also Figure S8A for data from individual brains. **C.** Heatmap showing normalized, averaged ERK intensity of the same brains as in A. Log2 fold changes (triploid/diploid) of ERK intensity along Y axis are plotted to the right. Scale bar, 100 μm. **D.** Comparison of ERK intensity in the indicated brain regions. Each dot represents the mean value of one clutch. Grey lines connect dots from the same clutch. Crossbars denote mean ±SEM. ns, not significant, paired t test. See also Figure S8B for ERK RNA-seq results.

### Triploid tadpoles show distinct swimming behavior compared to diploids

We next asked whether physiological differences at the brain level translated to behavioral differences at the organism level by comparing swimming behavior of diploid and triploid tadpoles (Figures 6A and S9A). Diploid and triploid siblings were placed into side-by-side arenas and their swimming activity was recorded following acute vibration and analyzed manually. Tadpoles were binned into three categories based on how long they stayed active during the 2-minute recording. “Still” was defined as stationary or only showing very brief and unsustained swimming; “half active” was defined as showing sustained swimming for half of the time; and “active” was defined as swimming continuously throughout the recorded session (Figure 6B). For both ploidies, most tadpoles did not begin to move until 4 days post-fertilization (dpf) (stage 42-46), when the percentage of half-active tadpoles peaked. Approximately 80% of the half active tadpoles were active during the first half of the video (Figure S9B), indicating that tadpoles at this stage swam in response to stimulation, rather than spontaneously. From 5 dpf (stage 46-47) onwards, most tadpoles remained active throughout the entire session. These observations are consistent with previous studies showing that although the relevant sensory and motor systems to support simple swimming episodes are present by stages 37-39, tadpoles remain mostly stationary until stage 44-45^41–44^; while sustained spontaneous swimming likely involves the central nervous system and occurs later in development^43^. Thus, swimming behavior of tadpoles of both ploidies follows a similar developmental trajectory.

**Figure 6.**
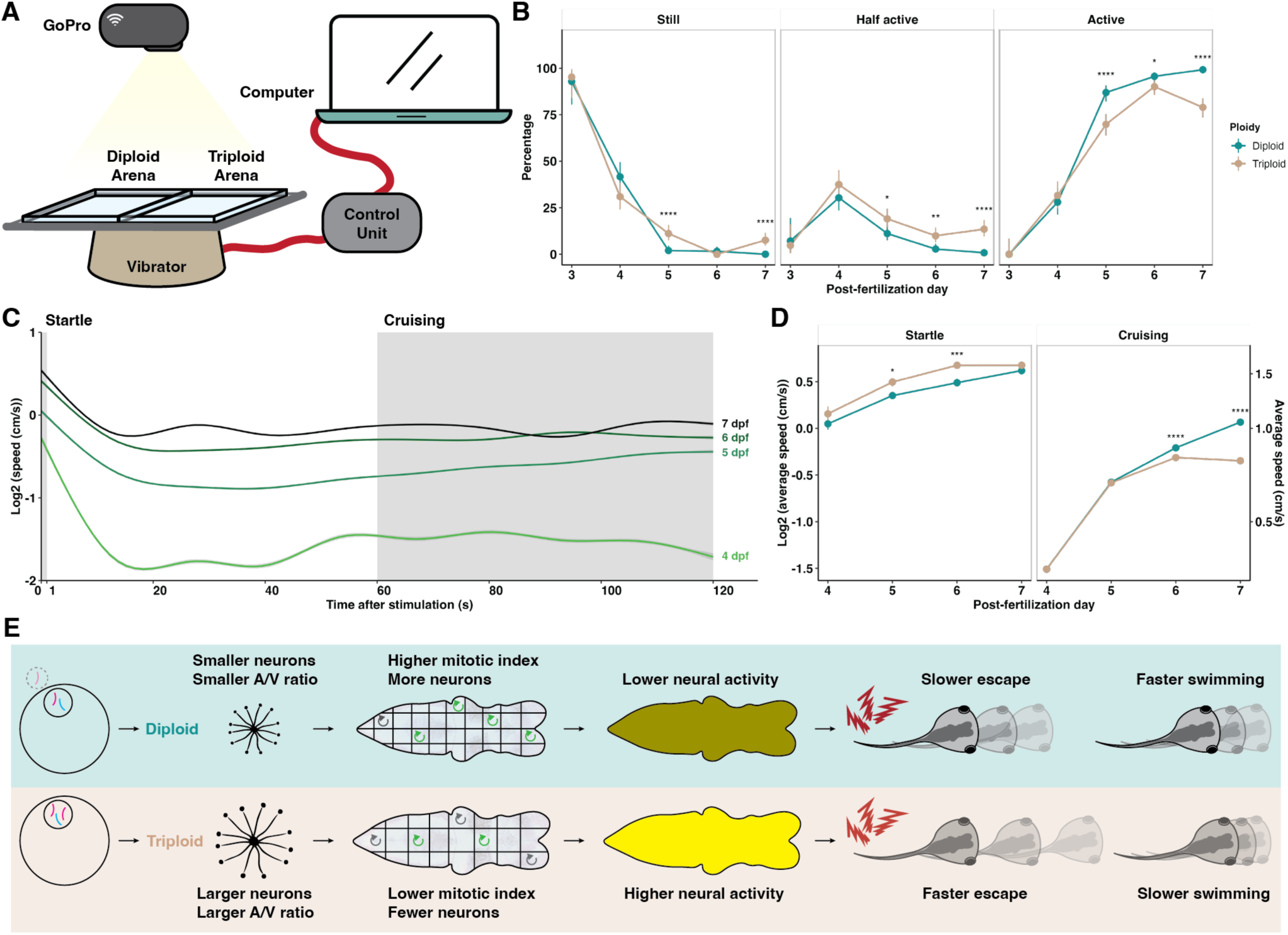
Neuron size and brain physiology differences produce distinct swimming behaviors in triploid tadpoles. **A.** Diagram showing the experimental setup of the swimming assay. See also Figure S9A for the time scheme and number of animals and replicates used. **B.** Manually scored categorizations of the tadpoles based on their activity level. Error bars mark 95% confidence interval and data points are connected to show the trend over development. *, p<0.05; **, p<0.01; ****, p<0.0001, Fisher’s exact test. See also Figure S9B for the breakdown of the “half active” category. **C.** Real-time speed of Trex-tracked swimming tadpoles during the recorded 2-minute window following stimulation. Diploid and triploid data were pooled. Data were smoothed with a generalized additive model (GAM) and presented as geometric mean ± 95% confidence interval. Grey boxes mark the time windows used to define the two different modes of swimming in D. **D.** Comparison of diploid and triploid speeds under the indicated swimming mode. Data were log2-transformed to correct for non-normality and are presented as geometric mean ± SEM. *, p<0.05; ***, p<0.001; ****, p<0.0001, t test. See also Figure S9C. **E.** Schematic summarizing results of this study.

As the tadpoles became increasingly active after 4 dpf, a difference between diploid and triploid behavior emerged. A significantly lower percentage of triploid tadpoles stayed active throughout the recorded session compared to their diploid siblings on 5, 6, and 7 dpf (Figure 6B, right panel). Conversely, a significantly higher percentage of triploid tadpoles remained still for half or all of the recorded durations (Figure 6B, left and middle panels).

To determine whether triploid tadpoles swam less because their swimming behavior or apparatus was less developed or defective, we utilized the open-source algorithm TRex^45^ and tracked real-time swimming speeds of individual tadpoles. As expected, tadpoles swam faster as they developed (Figure 6C, compare curves). By tracking tadpole speed over the 2-minute time span following stimulation, we identified two distinct swimming modes (Figure 6C, compare within each individual curves). Tadpoles shifted from a faster speed in the first few seconds following stimulation to a slower cruising mode when no further stimulation was applied. We denoted the first second after stimulation as the “startle” time window and the last 60 seconds as the “cruising” time window (Figure 6C, grey boxes), and calculated the average swimming speed of tadpoles in these two windows. Compared to their diploid siblings, triploid tadpoles swam significantly faster during their startle response, but significantly slower when in cruising mode (Figure 6D). This distinct swimming pattern indicates that motor circuit wiring is impacted by neuron size. Moreover, the observation that triploids could reach a faster swimming speed compared to diploids when startled suggests that the swimming difference is likely not due to muscle deficits.

With this swimming assay, we also examined whether repeated stimulation induced distinct learning patterns between diploid and triploid tadpoles. We did not observe differences in either startle or cruising speed across 7 repetitions in one experimental session in either ploidy (Figure S9C).

Altogether, our data indicate that differences in neuron size between the two ploidies lead to altered cell cycle dynamics at the cellular level, changed physiology and activity at the brain/organ level, and distinct behavioral patterns at the organism level, identifying a significant impact of cell size on the development and function of the vertebrate nervous system (Figure 6E).

## DISCUSSION

### Neuron size beyond cell body size

Previous studies on neuron size to genome size scaling measured the cell body^7,8^. Neurite size is important in determining neuronal activity and connectivity, and cannot be adequately represented by cell body size^13,46^. By profiling individual neuron size and shape *in vivo*, this work provides new insight into how different parameters of neuron size scale with genome size in the brain of a vertebrate. We show that neuron volume scales linearly with genome content, with neurons maintaining their general structure and neurite to cell body volume ratio. This isometric scaling likely reflects intrinsic needs and regulatory mechanisms that allocate and balance volume and cellular resources between compartments to ensure operation of basic, housekeeping cellular functions while achieving optimal information processing. Interestingly, the increase in neurite volume in triploid neurons is driven entirely by increased length, with a slight downward shift in triploid neurite diameter. Neurite growth requires microtubule motor-based transport and the degree of microtubule bundling has been shown to affect the diameter and extension rate of neurites^47,48^. Fast growing axons are often “stretched” and correlate with smaller diameters^47,49^. It is therefore possible that the smaller diameter of triploid neurites reflects faster extension as they grew almost twice as much in length as diploid neurites during the 3 days following labeling. Future studies could compare neurite growth rate between ploidies and examine whether triploid neurites catch up in diameter later in development.

A direct consequence of decreased neurite diameter is the increase in surface area to volume ratio. We report a 1.65-fold increase in total surface area in triploid neurons compared to diploids, superlinear to the ploidy and volume increase. This is likely an underestimate because only smaller neurons could be profiled. The larger the neuron, the higher proportion of both the volume and surface area is contributed by the neurite compartment (Figure S3). Therefore, surface area to volume ratio of a neuron would increasingly lean towards that of its neurite compartment as neuron size increases. We estimate that for larger neurons, the increase in surface area would approach 1.71-fold, that of the neurite compartment. Superlinear scaling of neuronal surface area has multiple physiological implications. The expansion of surface area in relation to volume alleviates constraints on the number of channels, pumps, signaling molecules, and synaptic machinery that can fit on a limited membrane area^52^, impacting membrane potential control^50^ as well as crucial neurodevelopmental processes including axon guidance^51^, target recognition^52^, and synapse formation^53^. On the other hand, an increase in surface area could place a heavier burden on cellular biosynthesis and energy consumption^27,54,55^ in triploid brains.

### Triploid brains show a slight but significant increase in size

Organ size is determined by both the size and number of constituent cells. We report that in the brains of *X. laevis* triploids, the increase in neuron size is largely offset by a decrease in neuron number, resulting in a small increase in overall brain size compared to diploids. In accordance with this, we also report a lower mitotic index in triploid brains. Whether effects on the cell cycle are autonomous or regulated as part of an active mechanism sensing and modulating brain size remains a fascinating question. Interestingly, brain size scales linearly between *X. laevis* and *X*. *tropicalis* (compare Figure 3D and S4G), corresponding to differences in genome and cell size^27^. The disparity in how brain size responds to changes in cell size in acute versus evolutionary ploidy changes could reflect different mechanisms of organ size control. Early amphibian limb graft experiments revealed that intrinsic and extrinsic mechanisms co-exist in vertebrate organ size control^56,57^. It would be interesting to perform neural tube grafts between ploidies and species and observe how brain size homeostasis operates with altered extrinsic signals.

It is worth noting that the increase in brain size following the ploidy and neuron size increase is significant and non-negligible in both *X. laevis* and *X. tropicalis*. This could be a coincidence if the lowered proliferation and neuron count in triploid brains is regulated in an entirely autonomous manner. On the other hand, that triploid brains do not grow to the exact same size as diploid brains could reflect possible physical and/or wiring constraints. For example, the increased total surface area in triploid brains could result in a larger volume of extracellular space (ECS), increasing total brain volume on top of the volume occupied by brain cells. Previous studies using *in vivo* diffusion analyses in rats and theoretical modeling have reported that ECS averages 20-60 nm in width and makes up about 20% of total brain volume^58–60^. As ECS volume scales positively with the total surface area of brain cells, it is likely that triploid brains contain a larger ECS compared to diploid brains. Moreover, the nervous system relies on intricate wiring for both development and function and in some cases the number of essential neurons cannot simply be scaled down. An extreme example is the Mauthner cells, a pair of giant neurons located in each side of the hindbrain involved in escape behaviors^61^. Both diploid and triploid tadpoles have one pair of Mauthner cells, and the connections of these neurons at the brain and spinal cord levels do not allow triploid brains to possess fewer than two. Our cell counting methods, which assess the whole brain or one brain region in bulk and provide reliable average values, would not reveal subtle and smaller-scale deviations from an inverse-linear scaling relationship.

### Ploidy and neuron size changes impact nervous system function

The drastic decrease in neuron numbers in triploid brains predicts changes in the wiring pattern and neuronal activity. We used phosphorylation of ERK, a downstream event of calcium influx, as a reporter to examine neuronal activity^38^. Traditionally, especially in mammals, expression of immediate early genes (IEG) like c-Fos is used to report neuronal activity^62^. However, c-Fos was not suited for our study because of its low sensitivity^63^, variability among brain regions^62,64^, low baseline staining in fish^39^ and amphibians^65^, and practically, lack of robust antibody staining compatible with optical clearing. Upstream of IEG^66^, pERK is a reliable marker for active neurons^39,40,67–71^. Its higher expression allows assessment of baseline neuronal activity in unstimulated brains and co-staining with total ERK provides a control to normalize staining variability.

We report a global increase in neural activity in triploid brains as read out by pERK/ERK levels. This is consistent with our model that a greater membrane surface area to volume ratio changes the number and distribution of ion channels, altering membrane potential control and downstream cellular responses like ERK phosphorylation. However, while our data show that pERK levels respond to extrinsic stimuli triggering neuronal activity, future studies utilizing electrophysiology^72^ are needed to further explore the electrical properties of diploid and triploid neurons.

At the organism level, we show that triploid tadpoles are more dormant and swim at a slower speed than their diploid siblings when unstimulated. Interestingly, a previous study has reported that polyploid *Ambystoma* salamanders walk a shorter distance on treadmill trials than diploids, making them inferior dispersers in nature^73^. Although neuron size and neuronal activity was not explored in that study, similarity with our results is unlikely to be a coincidence given the general positive scaling relationship between neuron size and genome size in amphibian species^7^.

The slower swimming speed of triploid tadpoles is likely not due to muscle deficits, since the animals were capable of sustained swimming for more than a few seconds at higher speeds. In fact, we show that triploid tadpoles swim significantly faster than diploids when stimulated by vibration. The response to hydrodynamic disturbance involves mechanosensory lateral line system as well as motor nuclei within the hindbrain^42^. How the difference in vibration response between ploidies arises remains an intriguing question. For future studies, calcium imaging^74^ could be utilized to assess neuronal activation before and after stimulation. Additionally, it would be interesting to test tadpole responses to various stimuli involving different sensory systems located in different brain regions.

Our findings highlight the functional consequences of genome and neuron size changes at both the organ and whole animal level. The interplay between genome size, cell size, and cellular physiology is complex. We found little alteration in gene expression patterns between diploid and triploid brains that could account for the physiological differences observed. The similar transcriptional landscape between ploidies is not unexpected. Previous studies in yeast and mammalian cells have shown that transcription is buffered against DNA dosage and that mechanisms are in place to measure the ratio of cellular volume to DNA content^31,75,76^. In the *C. elegans* intestine, transcription is sensitive to cellular volume to DNA ratio rather than to ploidy^77^. In zebrafish, comparisons among haploid, diploid, and tetraploid embryos revealed no significant transcriptional alterations across ploidies^78^. Moreover, studies in plants show that autopolyploids generally display lower alteration of gene expression due to the uniform increase in genome content, and that transcription might respond to genome content change over many generations following the initial polyploidization event^79–81^. Therefore, instead of the ploidy change giving rise to transcriptional disparities, we propose that the altered neural development and function in triploid embryos stems from post-transcriptional regulation in response to changes in neuron size and shape. Taken together, our work provides a framework and a tractable model for future research to further elucidate the cellular and molecular links between neuron size and neurodevelopment.

## Acknowledgements

We thank Marla Feller, Helen Bateup, Meng-meng Fu, Marc Hammarlund, Coral Zhou, Helena Cantwell, Gabriel Cavin-Meza, Kevin Williams, and Shenliang Yu for feedback on the manuscript, Helen Willsey, Kate McCluskey, Na Ji, and Gokul Upadhyayula for discussion and suggestions on reagents and methodology, Caroline McKeown (Hollis Cline lab, Scripps) for advice on electroporation, Shaina Carroll and Kartoosh Heydari for help with flow cytometry experiments, Virgilio López III (Carlos Aizenman lab, Brown University) for advice on tadpole swimming assays, and Martin Ziyuan Liu and the animal facility of UC Berkeley for frog care. R.H. was supported by NIH MIRA grant R35GM118183652 and the Flora Lamson Hewlett Chair in Biochemistry.

## AUTHOR CONTRIBUTIONS

Conceptualization, X.L. and R.H.; Methodology, X.L.; Investigation, X.L., C.W., and S.A.S.; Data Curation, X.L., C.W., and S.A.S.; Formal analysis, X.L.; Visualization, X.L.; Writing – Original Draft, X.L.; Writing – Review & Editing, X.L. and R.H.; Project Administration: X.L.; Funding acquisition, R.H.; Supervision, R.H.

## STAR⍰METHODS

### Frog care

All frogs were used and maintained following standard protocols established by the UC Berkeley Animal Care and Use Committee and approved in our Animal Use Protocol. Mature *X. laevis* and *X. tropicalis* were obtained from Nasco (Fort Atkinson, WI) or the National *Xenopus* Resource Center (Woods Hole, MA), and were housed in a recirculating tank system with regularly monitored temperature and water quality, with *X. laevis* at 20-23°C and *X. tropicalis* at 23-26°C. All animals were fed Nasco frog brittle.

*X. laevis* and *X. tropicalis* females were ovulated with no harm to the animals with a minimum 4-months rest interval. To obtain testes for *in vitro* fertilizations, *X. laevis* and *X. tropicalis* males were euthanized by over-anesthesia through immersion in 0.05% benzocaine in double-distilled water (ddH_2_O) prior to dissection. Carcasses were frozen at -20°C. Tadpoles were euthanized in 0.01‰ benzocaine in ddH_2_O prior to fixation.

### In vitro fertilization of X. laevis and X. tropicalis

*Xenopus in vitro* fertilizations were performed as previously described^29^. *X. laevis* females were primed with 100 U of pregnant mare serum gonadotropin (PMSG) (Calbiochem #367222) >48 h before boosting, boosted with 500 U of human Chorionic Gonadotropin (hCG) (either Sigma #CG10, or Chorulon Merck #133754) the afternoon before the experiment, and kept at 16°C overnight in 1X MMR (100 mM NaCl, 2 mM KCl, 2 mM CaCl_2_, 1 mM MgSO_4_, 0.1 mM EDTA, 5 mM HEPES-NaOH, pH 7.6). *X. tropicalis* females were primed with 10 U hCG 14-16 h before boosting, kept at room temperature in ddH_2_O, and boosted with 250 U the morning of the experiment. Testes were dissected from euthanized males and kept at 4°C in 1X MR (100 mM NaCl, 1.8 mM KCl, 1 mM MgCl_2_, 5 mM HEPES-NaOH, pH 7.6) for *X. laevis* or placed in 1X MBS (88 mM NaCl, 1.006 mM KCl, 2.49 mM NaHCO_3_, 0.998 mM MgSO_4_, 5 mM HEPES-NaOH, pH 7.8) with 0.2% Bovine Serum Albumin (BSA) and used the same day for *X. tropicalis*.

Eggs were collected by gently squeezing females atop a 60 mm petri dish. Sperm solution was prepared by homogenizing a small piece of testis (∼1/4 for *X. laevis*, ∼1/2 for *X. tropicalis*) in 1 mL of 1/10X MMR (*X. laevis*) or 0.5 mL of 0.2% BSA in 1X MBS *(X. tropicalis)* in a 1.7 mL Eppendorf tube using a plastic pestle. Eggs were fertilized with the sperm solution, gently swirled until they formed a monolayer at the bottom of the dish, incubated for 10 min (*X. laevis*) or 4 min *(X. tropicalis)* with the dish slightly tilted to ensure submersion of eggs, and then flooded with 1/10X MMR (*X. laevis*) or ddH_2_O *(X. tropicalis)*.

To obtain triploid embryos, pairs of dishes were fertilized. Control diploid dishes were kept in 1/10X MMR until de-jellying. Triploid dishes were cold shocked at 13 min (*X. laevis*) or 9 min *(X. tropicalis)* post fertilization with pre-chilled 4°C 1/10X MMR in a slushy ice bath for 15 min. After the cold shock, embryos were equilibrated in room temperature 1/10X MMR for at least 5 min. To remove jelly coats, embryos were incubated in freshly prepared De-jellying Solution (3% L-cysteine in ddH2O-NaOH, pH 7.8) for about 10 min, and then washed 5X with 1/10X MMR.

### Maintenance and staging of *Xenopus* embryos

Fertilized and de-jellied embryos were transferred to larger dishes (100 mm diameter X 15 mm high) and kept in 1/10X MMR in incubators set at 23°C (for *X. laevis*) or 24°C (for *X. tropicalis*). Dead or lysed embryos were removed, and media was exchanged with fresh 1/10X MMR twice a day for the first two days and at least once a day afterwards. Diploid and triploid embryos were maintained at the same density and media levels.

Tadpoles were euthanized and staged before experiments according to the Nieuwkoop and Faber development table^84^.

### Tadpole sex determination

PCR-based sex determination was performed as previously described^27,85^. Tails of individual euthanized tadpoles were cut, transferred to 20 μL of 1X Phusion buffer (New England Biolabs #E0553S) in 200 μL PCR tubes, frozen at -80°C for >1 h, thawed, spun down, and boiled at 95°C for 10 min. Next, 2.5 μL of 20 ng/mL Proteinase K (New England Biolabs #P8107S) was added to each tube and the tubes were incubated at 55°C for 4 h, boiled at 95°C for 10 min, and stored at 4°C. 2.5 μL of the lysis solution was used as template in the subsequent PCR reaction, which was performed using the Phusion High-Fidelity PCR kit (New England Biolabs E0553S), forward (AAAACCATGACCTCCCGGATAC) and reverse (TAGGGAGGGGTTTGGAGGTTC) primers^85^, with 35 cycles annealing at 58°C for 30 s and elongating at 72°C for 30 s.

### Tadpole brain immunofluorescence and clearing

Immunofluorescence and clearing was performed combining protocols from Helen Willsey’s lab (UCSF, CA) and Affaticati et al^86^. Euthanized tadpoles were fixed in 4% PFA in PBS at 4°C overnight and washed 4X with PBS. For *X. laevis*, 5-10 brains of staged tadpoles were dissected using two pairs of fine-tip tweezers and collected in 500 μL of PBS in a 1.7 mL Eppendorf tube. For *X. tropicalis*, 5-7 tadpoles were directly collected without brain dissection.

To bleach the samples, 500 μL of 2X Bleaching Solution (1X: 10% formamide, 8% hydrogen peroxide in PBS) was added to each tube and the tubes were placed in a light-colored rack under bright light for 1.5 h. Bleached samples were washed extensively (>4X) with PBST, incubated in fresh PBST overnight at 4°C, blocked and permeabilized with 50 μL of Blocking Buffer (10% goat serum (Jackson ImmunoResearch #005-000-001), 10% DMSO, and 1% Triton X-100 in PBST) for 2-4 h at room temperature, and then incubated with primary antibodies in 50 μL of Staining Buffer (2% goat serum, 10% DMSO, 0.1% Triton X-100, and 0.05% sodium azide in PBST) for 2-3 days at 4°C. Next, samples were washed three times each for >20 min with PBST at room temperature and incubated with secondary antibodies along with 10 ng/mL Hoechst 33342 (Thermo Fisher Scientific #H3570) in 50 μL of Staining Buffer for 1-2 days at 4°C, with the tubes wrapped in foil to avoid light. All incubations were carried out on a nutator. All primary and secondary antibodies were used at 1:250 including mouse anti-HuC/HuD (Thermo Fisher Scientific #A-21271), mouse anti-β-tubulin (DSHB E7 concentrate form), mouse anti-PCNA (Thermo Fisher Scientific #13-3900), mouse anti-p44/42 MAPK Erk1/2 (Cell Signaling #4696), rabbit anti-phospho-p44/42 MAPK Erk1/2 (Thr202/Tyr204) (Cell Signaling #4370), rabbit anti-phospho-Histone H3 (Ser10) (Millipore Sigma #06-570), rabbit anti-RFP (Abcam #ab62341), rabbit anti-cleaved caspase-3 (Cell Signaling #9661), Alexa Fluor 488 goat anti-mouse IgG (Thermo Fisher Scientific #A-11001), Alexa Fluor 568 goat anti-mouse IgG (Thermo Fisher Scientific #A-11004), Alexa Fluor 488 goat anti-rabbit IgG (Thermo Fisher Scientific #A-11008), and Alexa Fluor 568 goat anti-rabbit IgG (Thermo Fisher Scientific #A-11011).

To clear samples after the immunostaining steps, fructose-based high-refractive index solution (fbHRI) was prepared by mixing 4 parts of F1457 (90% fructose in fbHRI diluent (0.2X PBS, 0.002% sodium azide), refractive index (RI) 1.457) with 1 part of DMSO1457 (85% DMSO in fbHRI diluent, RI 1.457) and adjusting RI to 1.457 with 100% fructose/fbHRI diluent. Stained samples were washed three times each for >20 min with PBST at room temperature on a nutator, incubated for >2 h in PBST, exchanged into 50% fbHRI in fbHRI diluent, incubated for >6 h at 4°C, exchanged into fbHRI, and incubated for >12 h at 4°C. Cleared samples were stored in fbHRI, avoiding light, at 4°C until imaging, for up to a few weeks.

### *In toto* imaging of whole-mount brains

Stained and cleared brains (for *X. laevis*) or tadpoles (for *X. tropicalis)* were mounted in fbHRI onto small glass bottom dishes (MatTek # P35G-1.5-14-C), with the dorsal side facing down. Extra liquid was removed so that the brain would lay flat on the glass bottom. Confocal imaging was performed on an inverted ZEISS LSM 800 using the ZEISS ZEN 2.3 (Blue edition) software, with a Plan-Apochromat 10X/0.45 air objective (for brain size and morphology measurements), a Plan-Apochromat 20X/0.8 air objective (for pERK/ERK intensity measurements), or a Plan-Apochromat 40x/1.2 Imm Corr DIC glycerine immersion objective (for nucleus counting, nuclear volume measurements, and single neuron imaging), with a pinhole size set at 1 Airy Unit. In experiments quantifying and comparing fluorescence intensity, all laser settings were kept the same for diploid and triploid samples and images were taken in 16-bit. Post-hoc brightness/contrast adjustments, if any, were always applied uniformly across diploid and triploid samples. In experiments where fluorescence intensity was not quantified, laser settings were adjusted separately utilizing the Range Indicator Tool in the ZEN software for optimal visualization, and brightness/contract adjustments may differ between samples for presentation purposes. Image stitching, when needed, was performed either with the built-in Tile Function of the ZEN software, or the Stitching Plugin^87^ of the FIJI software^88^.

### Sparse labeling and single neuron reconstruction

Electroporation was performed mostly following protocols from Hollis Cline’s lab^32,33^. On 4 dpf, stage 44 tadpoles were anesthetized in 0.01‰ benzocaine in 1/10X MMR for several minutes and transferred to a wet tissue on top of an inverted petri dish. 20-30 nL of Injection Mix (350-400 ng/μL pCMV-RFP plasmid, 0.2% Brilliant Blue dye in EB buffer) was injected into the brain ventricle of each tadpole using needles pulled from glass capillaries (World Precision Instruments #TW100F-4), mounted on a Narishige micromanipulator, and connected to a Picospritzer III Microinjection Dispense System. Next, a pair of platinum electrodes, cut from Sutter puller filaments, controlled by another Narishige micromanipulator, and connected to a Grass SD9 Stimulator, were positioned to clamp the desired brain region of the injected tadpole. Two 30V, 1.6 ms pulses each polarity were delivered through the electrodes.

Electroporated tadpoles were transferred to large dishes with fresh 1/10X MMR in a well-ventilated area for recovery. When movement was observed again, tadpoles were put back into 23°C incubators and allowed to rest for 3 days. On 7 dpf, tadpoles were euthanized and fixed, and their brains dissected, stained with anti-RFP and anti-HuC/D antibodies, cleared, and imaged as described earlier. Diploid and triploid samples to be compared were always processed in parallel through the injection, electroporation, recovery and subsequent staining and imaging steps.

Labelled single cells with confirmed neuronal identity (HuC/D positive) were reconstructed with Imaris 10.2.0 (license owned by CRL Molecular Imaging Center, UC Berkeley) semi-automatically based on the immunostaining-amplified RFP signal, with the Surfaces Function to remove background signal and the Filaments Function to reconstruct the soma model and neurite paths (with soma as starting point and no loops). Parameters and settings in automatic programs were kept constant between all processed neurons when possible, and manual supervision and model training were performed with best judgement. After reconstruction, all measurements were exported and analyzed with R 4.2.1^89^.

### Imaging-based brain measurements

For nucleus counting and nuclear volume measurements, brains were dissected, stained with anti-PCNA and anti-caspase-3 antibodies and Hoechst, and cleared as described earlier. Whole-mount brains were imaged with confocal microscopy as described earlier except that the Auto Z Brightness Correction Function in the ZEN software was used to compensate for dimmer fluorescence in deeper tissues. Two 159.73 μm X 159.73 μm X 100 μm regions of interest (ROIs) in left and right forebrain and a 319.45 μm X 319.45 μm X 50 μm ROI in hindbrain were cropped from the image stack of each brain. Similar regions were cropped across all processed brains using morphological landmarks as reference. ROI was not taken in midbrain because light refraction across the midbrain ventricle caused images to be less sharp. Single nuclei were segmented with Cellpose^90^ using the Cyto Model based on the Hoechst channel. Segmented nuclei were counted and measured using the 3DSuite Plugin^91^ of the FIJI software^88^. A home-built code in R 4.2.1^89^ was used to remove measurements of nuclei positive for PCNA (progenitors, not neurons) or at ROI borders. Caspase positive nuclei were counted manually because the labelling was sparse.

For brain morphology measurements, dissected brains (for *X. laevis*) or tadpoles (for *X. tropicalis)* were stained with anti-β-tubulin antibody and Hoechst and imaged as described earlier except that to avoid potential shrinkage, samples were not cleared and were mounted and imaged in PBST. Forebrain, midbrain, and hindbrain ROIs were traced manually using the Freehand Selections Tool in FIJI^88^. Sample identity was masked during this process to minimize bias. All measurements were exported and analyzed with R 4.2.1^89^.

### Flow cytometry and cell cycle analysis

Dissected brains from tadpoles of the correct stage were processed for staining (with Hoechst, with or without anti-PCNA and anti-PH3 antibodies) as described above without the clearing steps. To dissociate the stained brains, 5-6 brains were transferred to 100 μL of Dissociation Buffer (90% StemPro Accutase (Thermo Fisher Scientific #A1110501), 0.05% Triton X-100, in PBS) in a 1.7 mL low retention Eppendorf tube and incubated at room temperature, avoiding light, overnight. The Accutase reaction was stopped by adding 250 μL of PBS to the tube and the brains were further physically dissociated by pipetting up and down for 30 s. 20 μL of CountBright Absolute Counting Beads (Thermo Fisher Scientific #C36950, LOT #2361081, 0.52*10^5^ beads/50 μL) were added to each tube, mixed with the solution by pipetting up and down 10 times, and the whole mixture was transferred to a 12 mm X 75 mm CellPro flow cytometry tube through a 35 μm mesh strainer cap (Alkali Scientific #CT6405). The Eppendorf tube was washed twice each with 250 μL of 1X HBSS, and the washing liquid was also transferred through the mesh to the flow cytometry tube. Flow cytometry tubes were kept on ice, avoiding light, until the flow cytometry run. Low retention pipet tips were used throughout.

Each tube of dissociated brain cells was vortex briefly and run with a BD LSR Fortessa X-20 Cell Analyzer (CRL Flow Cytometry Facility, UC Berkeley). Pacific Blue channel was used for detecting Hoechst staining, and FITC channel for Alexa Fluor 488, PE-Texas Red channel for Alexa Fluor 568, APC-cy7 channel for the rainbow-colored counting beads. A tube of dissociated brain cells with beads added but without any staining was included in each run to facilitate gate drawing. FlowJo 10.8.1 (Becton Dickinson & Company) was used for processing the flow cytometry data. All events were gated with forward scatter width over forward scatter area (FSC-W/FSC-A) for singlets. Singlets were separated into a “Beads” and a “Non-beads” population based on APC-cy7 channel intensity. The “Non-beads” population was further gated first with side scatter width over side scatter area (SSC-W/SSC-A) and then Hoechst intensity to ensure that they were single cell events. Next, for any one sample to pass the post-hoc quality control (QC) performed with a home-built code in R 4.2.1^89^, the following conditions were met: a) sample had > 5,000 bead events; b) sample had >10,000 single cell events; c) sample mean of FSC-A and Pacific Blue channel intensity of bead events fell inside run mean*(1±CV) of all bead events; d) sample CV of FSC-A and Pacific Blue channel intensity of bead events was less than 2X run CV of all bead events; and e) sample had a corresponding diploid/triploid sample to compare to. These gating and QC steps were performed in an unbiased way across all diploid and triploid samples in each run.

For cell counting, number of cells per brain (*N*) was calculated as:

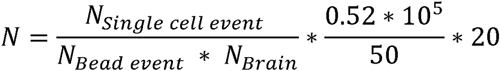

For initial cell cycle analysis (Figure 4A), FlowJo built-in Cell Cycle Tool was used on single cell populations with the following settings: Pacific Blue channel (Hoechst), Dean Jett Fox (DJF) model^83^, no synchronized peak, no peak constraints. For further cell cycle analysis combining PCNA and pH3 staining (Figures 4E-G and S5), a home-built R 4.2.1^89^ code was used to fit Gaussian peaks under probability density curves of logarithmic PCNA and pH3 intensities. Areas of each fitted peak were calculated as fractions of the corresponding population (Figures 4E-F).

### pERK/ERK intensity measurements

To measure baseline neural activity, unstimulated, freely swimming tadpoles were over-anesthetized in 0.05% benzocaine for 1 min and immediately processed for fixation. Corning Netwell Strainers were used to enable efficient buffer exchanges to minimize disturbance of tadpoles and potential impact on neural activity. Brains of stage 46 tadpoles were dissected, stained with anti-ERK and anti-pERK antibodies and Hoechst, cleared, and imaged as described earlier. Tiled images of each brain were auto-stitched into a 640.15 μm X 1790.04 μm z-stack in ZEISS ZEN and then analyzed with FIJI^88^. 170 z-slices across 150 μm from the top surface of the brain were summarized to a z-projection and registered to one reference brain projection generated from a representative, normal-looking stage 46 diploid brain using the bUnwarpJ Plugin^92^. With the registered images, pERK intensity was normalized to ERK intensity using the Image Calculator Function, and the normalized result was masked to remove background signal outside the brain sample area with a mask generated by thresholding the ERK channel with the “Huang” method followed by “Process>Binary>Fill holes”. ROIs of different brain regions manually traced with the Freehand Selections Tool on the reference projection were used for intensity measurements. To adjust for clutch variance, diploid images of each clutch were averaged, and the maximum intensity of the averaged image was used to normalize all measurements of that clutch. The summary images presented (Figures 5 and S7D) were averaged from clutch averages.

To measure neural activity after stimulation, all procedures were the same as above except that prior to euthanasia, tadpoles were split evenly between a tapping dish and a control dish, with the former manually tapped on bench top continuously for 15 min, and the latter placed on the farthest bench in the same room to avoid disturbance from the tapping noise. Maximum intensity of control averages of each clutch was used to normalize against clutch variance. Control data in this set of experiment (3 clutches, Figure S7D) were also included as diploid data in the unstimulated dataset (6 clutches, Figure 5).

### Brain-specific RNA-seq

J strain (37 generations inbred) *X. laevis* obtained from National *Xenopus* Resource were used to prepare samples for RNA-seq experiments. In the afternoon on 4 dpf, tadpoles were over-anesthetized in 0.05% benzocaine in 1/10X MMR for 1 min and immediately processed for dissection. For each replicate, >10 brains per ploidy of stage 44-46 tadpoles from the same clutch were dissected using two pairs of fine-tip tweezers and immediately transferred to a 1.7 mL Eppendorf tube containing 100 μL of PBS and 400 μL of TRIzol (Invitrogen #15596026). Tubes containing dissected brains were kept on ice during dissection, fast frozen with liquid N_2_ immediately after dissection, and kept at -80°C.

To extract RNA, brains were subjected to 3 cycles of freeze-thaw with liquid N_2_ and a 37°C water bath and incubated at room temperature for 5 min. Next, 80 μL of chloroform was added to each tube and the mixture was inverted several times for 15 s, incubated at room temperature for 3 min, transferred to a Phasemaker tube (Invitrogen # A33248), and spun at 12,000 g at 4°C for 15 min. The aqueous phase was transferred to a new 1.7 mL Eppendorf tube, mixed with 200 μL of isopropanol, incubated at room temperature for 10 min, and spun at 12,000 g at 4°C for 10 min. After carefully removing the supernatant, the pallet was washed with 400 μL of pre-chilled 75% ethanol twice, air-dried for 5 min on ice, and resuspended in 44 μL of RNase-free water. Potential DNA contaminant was removed by incubating with 5 μL of 10X DNase I buffer and 1 μL of DNase I (NEB #M0303S) at 37°C for 10 min, and RNA was purified using the Monarch RNA Cleanup Kit (NEB #T2040S).

RNA quality control, library preparation (Kapa Biosystems library preparation kits with covaris/bioruptor shearing for gDNA, custom Unique Dual Indexes to eliminate cross sample bleed), and sequencing (NovaSeq 6000 S4, 150 bp paired-end reads, 25M reads per sample) was performed by QB3 Genomics, UC Berkeley. Sequencing results were fastQC-ed^93^, quality trimmed with Trim Galore^94^, and aligned to reference transcriptome (*Xenopus laevis* v10.1) with Salmon^95^ in Linux. Data from Salmon was imported to DESeq2 using TXimport^96^ and differential expression analysis was performed with DESeq2^97^ in R 4.2.1^89^.

### Tadpole swimming assay

The tadpole swimming assay setup (Figure 6A) was inspired by a protocol from Lopez et al^98^. For each session, 6 tadpoles each ploidy from the same clutch were placed in 1/10X MMR in a pair of swimming arenas, made from 100 mm X 100 mm clear plastic dishes, arranged side by side, lighted from below with an LED screen, and attached to a computer-controlled dental vibrator. Each session consisted of 7 consecutive repeats, in each of which tadpoles were stimulated by a brief, strong vibration, recorded for 2 min by a GoPro camera fixed above the arenas, and allowed to rest without stimulation for another 3 min. Tadpoles were euthanized afterwards, and different groups of 6 tadpoles each ploidy were used for sessions at different time points. Tadpole swimming in the recorded videos was first scored manually based on activity level, and then traced automatically with the open-source algorithm TRex^45^. Next, measurements were exported and analyzed with R 4.2.1^89^. The detailed time scheme and number of replicates used are reported in Figure S9A.

## SUPPLEMENTAL INFORMATION TITLES AND LEGENDS

Document S1. Figures S1-S9

Figure S1 Polyploid *Xenopus* as a model to study brain development, related to Figure 1

Figure S2 Size and shape analysis of diploid and triploid neurons, related to Figure 1

Figure S3 Proportion of neuronal volume and surface area contributed by neurites, related to Figure 2

Figure S4 Triploid brains are morphologically similar to diploid brains, related to Figure 3

Figure S5 Triploid brains show distinct cell birth/death dynamics, related to Figure 4

Figure S6 Triploid and diploid brains possess similar transcriptional profiles, related to Figure 4

Figure S7 pERK expression is neuronal and reflects neural activity, related to Figure 5

Figure S8 Triploid brains show increased pERK/ERK level but unaltered ERK expression, related to Figure 5

Figure S9 Assessing stimulated swimming, spontaneous swimming, and learning behavior, related to Figure 6

## REFERENCES

1. Lloyd AC. The Regulation of Cell Size. Cell. 2013;154(6):1194–1205. doi:10.1016/j.cell.2013.08.053

2. Ginzberg MB, Kafri R, Kirschner M. On being the right (cell) size. Science. 2015;348(6236):1245075. doi:10.1126/science.1245075

3. Azevedo FAC, Carvalho LRB, Grinberg LT, et al. Equal numbers of neuronal and nonneuronal cells make the human brain an isometrically scaled-up primate brain. J Comp Neurol. 2009;513(5):532–541. doi:10.1002/cne.21974

4. Herculano-Houzel S. The glia/neuron ratio: how it varies uniformly across brain structures and species and what that means for brain physiology and evolution. Glia. 2014;62(9):1377–1391. doi:10.1002/glia.22683

5. Mota B, Herculano-Houzel S. All brains are made of this: a fundamental building block of brain matter with matching neuronal and glial masses. Front Neuroanat. 2014;8:127. doi:10.3389/fnana.2014.00127

6. Levy DL, Heald R. Biological Scaling Problems and Solutions in Amphibians. Cold Spring Harb Perspect Biol. 2016;8(1):a019166. doi:10.1101/cshperspect.a019166

7. Roth G, Walkowiak W. The Influence of Genome and Cell Size on Brain Morphology in Amphibians. Cold Spring Harb Perspect Biol. 2015;7(9):a019075. doi:10.1101/cshperspect.a019075

8. Roth G, Blanke J, Wake DB. Cell size predicts morphological complexity in the brains of frogs and salamanders. Proc Natl Acad Sci. 1994;91(11):4796–4800. doi:10.1073/pnas.91.11.4796

9. Nandakumar S, Grushko O, Buttitta LA. Polyploidy in the adult Drosophila brain. VijayRaghavan K, Boekhoff-Falk GE, eds. eLife. 2020;9:e54385. doi:10.7554/eLife.54385

10. Nandakumar S, Rozich E, Buttitta L. Cell Cycle Re-entry in the Nervous System: From Polyploidy to Neurodegeneration. Front Cell Dev Biol. 2021;9:698661. doi:10.3389/fcell.2021.698661

11. Sigl-Glöckner J, Brecht M. Polyploidy and the Cellular and Areal Diversity of Rat Cortical Layer 5 Pyramidal Neurons. Cell Rep. 2017;20(11):2575–2583. doi:10.1016/j.celrep.2017.08.069

12. Yu L, Yu Y. Energy-efficient neural information processing in individual neurons and neuronal networks. J Neurosci Res. 2017;95(11):2253–2266. doi:10.1002/jnr.24131

13. Pannese E. III. Shape and Size of Neurons. In: Pannese E, ed. Neurocytology: Fine Structure of Neurons, Nerve Processes, and Neuroglial Cells. Springer International Publishing; 2015:13–23. doi:10.1007/978-3-319-06856-5_3

14. Goriounova NA, Heyer DB, Wilbers R, et al. Large and fast human pyramidal neurons associate with intelligence. Badre D, Behrens TE, Koch C, eds. eLife. 2018;7:e41714. doi:10.7554/eLife.41714

15. Kosillo P, Doig NM, Ahmed KM, et al. Tsc1-mTORC1 signaling controls striatal dopamine release and cognitive flexibility. Nat Commun. 2019;10(1):5426. doi:10.1038/s41467-019-13396-8

16. Kosillo P, Ahmed KM, Aisenberg EE, et al. Dopamine neuron morphology and output are differentially controlled by mTORC1 and mTORC2. eLife. 2022;11:e75398. doi:10.7554/eLife.75398

17. Bateup HS, Johnson CA, Denefrio CL, Saulnier JL, Kornacker K, Sabatini BL. Excitatory/inhibitory synaptic imbalance leads to hippocampal hyperexcitability in mouse models of tuberous sclerosis. Neuron. 2013;78(3):510–522. doi:10.1016/j.neuron.2013.03.017

18. Herculano-Houzel S, Manger PR, Kaas JH. Brain scaling in mammalian evolution as a consequence of concerted and mosaic changes in numbers of neurons and average neuronal cell size. Front Neuroanat. 2014;8:77. doi:10.3389/fnana.2014.00077

19. Rinne JO, Paljärvi L, Rinne UK. Neuronal size and density in the nucleus basalis of Meynert in Alzheimer’s disease. J Neurol Sci. 1987;79(1-2):67–76. doi:10.1016/0022-510x(87)90260-7

20. Rajkowska G, Selemon LD, Goldman-Rakic PS. Neuronal and glial somal size in the prefrontal cortex: a postmortem morphometric study of schizophrenia and Huntington disease. Arch Gen Psychiatry. 1998;55(3):215–224. doi:10.1001/archpsyc.55.3.215

21. Andrews CB, Gregory TR. Genome size is inversely correlated with relative brain size in parrots and cockatoos. Genome. 2009;52(3):261–267. doi:10.1139/G09-003

22. Gregory TR. Genome size and brain cell density in birds. Can J Zool. 2018;96(4):379–382. doi:10.1139/cjz-2016-0306

23. Backman SA, Stambolic V, Suzuki A, et al. Deletion of Pten in mouse brain causes seizures, ataxia and defects in soma size resembling Lhermitte-Duclos disease. Nat Genet. 2001;29(4):396–403. doi:10.1038/ng782

24. Kwon CH, Zhu X, Zhang J, et al. Pten regulates neuronal soma size: a mouse model of Lhermitte-Duclos disease. Nat Genet. 2001;29(4):404–411. doi:10.1038/ng781

25. Bassetti D, Luhmann HJ, Kirischuk S. Effects of Mutations in TSC Genes on Neurodevelopment and Synaptic Transmission. Int J Mol Sci. 2021;22(14):7273. doi:10.3390/ijms22147273

26. Exner CRT, Willsey HR. Xenopus leads the way: Frogs as a pioneering model to understand the human brain. Genes N Y N 2000. 2021;59(1-2):e23405. doi:10.1002/dvg.23405

27. Cadart C, Bartz J, Oaks G, Liu MZ, Heald R. Polyploidy in *Xenopus* lowers metabolic rate by decreasing total cell surface area. Curr Biol. 2023;33(9):1744–1752.e7. doi:10.1016/j.cub.2023.03.071

28. Brownlee C, Heald R. Importin α Partitioning to the Plasma Membrane Regulates Intracellular Scaling. Cell. 2019;176(4):805–815.e8. doi:10.1016/j.cell.2018.12.001

29. Gibeaux R, Heald R. Generation of Xenopus Haploid, Triploid, and Hybrid Embryos. Methods Mol Biol Clifton NJ. 2019;1920:303–315. doi:10.1007/978-1-4939-9009-2_18

30. Mueller RL. Genome Biology and the Evolution of Cell-Size Diversity. Cold Spring Harb Perspect Biol. 2015;7(11):a019125. doi:10.1101/cshperspect.a019125

31. Darmasaputra GS, van Rijnberk LM, Galli M. Functional consequences of somatic polyploidy in development. Dev Camb Engl. 2024;151(5):dev202392. doi:10.1242/dev.202392

32. Bestman JE, Cline HT. Morpholino Studies in Xenopus Brain Development. In: Sprecher SG, ed. Brain Development: Methods and Protocols. Humana Press; 2014:155–171. doi:10.1007/978-1-62703-655-9_11

33. Bestman JE, Lee-Osbourne J, Cline HT. In vivo time-lapse imaging of cell proliferation and differentiation in the optic tectum of Xenopus laevis tadpoles. J Comp Neurol. 2012;520(2):401–433. doi:10.1002/cne.22795

34. Schönenberger F, Deutzmann A, Ferrando-May E, Merhof D. Discrimination of cell cycle phases in PCNA-immunolabeled cells. BMC Bioinformatics. 2015;16:180. doi:10.1186/s12859-015-0618-9

35. Leonhardt H, Rahn HP, Weinzierl P, et al. Dynamics of DNA Replication Factories in Living Cells. J Cell Biol. 2000;149(2):271–280.

36. Prigent C, Dimitrov S. Phosphorylation of serine 10 in histone H3, what for? J Cell Sci. 2003;116(18):3677–3685. doi:10.1242/jcs.00735

37. Li DW, Yang Q, Chen JT, Zhou H, Liu RM, Huang XT. Dynamic distribution of Ser-10 phosphorylated histone H3 in cytoplasm of MCF-7 and CHO cells during mitosis. Cell Res. 2005;15(2):120–126. doi:10.1038/sj.cr.7290276

38. Rosen LB, Ginty DD, Weber MJ, Greenberg ME. Membrane depolarization and calcium influx stimulate MEK and MAP kinase via activation of Ras. Neuron. 1994;12(6):1207–1221. doi:10.1016/0896-6273(94)90438-3

39. Randlett O, Wee CL, Naumann EA, et al. Whole-brain activity mapping onto a zebrafish brain atlas. Nat Methods. 2015;12(11):1039–1046. doi:10.1038/nmeth.3581

40. Thyme SB, Pieper LM, Li EH, et al. Phenotypic Landscape of Schizophrenia-Associated Genes Defines Candidates and Their Shared Functions. Cell. 2019;177(2):478–491.e20. doi:10.1016/j.cell.2019.01.048

41. Sillar KT, Li WC. Neural control of swimming in hatchling *Xenopus* frog tadpoles. In: Whelan PJ, Sharples SA, eds. The Neural Control of Movement. Academic Press; 2020:153–174. doi:10.1016/B978-0-12-816477-8.00007-7

42. Saccomanno V, Love H, Sylvester A, Li WC. The early development and physiology of Xenopus laevis tadpole lateral line system. J Neurophysiol. 2021;126(5):1814–1830. doi:10.1152/jn.00618.2020

43. Roberts A, Li WC, Soffe SR. How neurons generate behavior in a hatchling amphibian tadpole: An outline. Front Behav Neurosci. 2010;4(JUN). doi:10.3389/fnbeh.2010.00016

44. Jamieson D, Roberts A. Responses of young Xenopus laevis tadpoles to light dimming: possible roles for the pineal eye. J Exp Biol. 2000;203(Pt 12):1857–1867. doi:10.1242/jeb.203.12.1857

45. Walter T, Couzin ID. TRex, a fast multi-animal tracking system with markerless identification, and 2D estimation of posture and visual fields. Lentink D, Rutz C, Pujades S, eds. eLife. 2021;10:e64000. doi:10.7554/eLife.64000

46. Brown KM, Gillette TA, Ascoli GA. Quantifying neuronal size: summing up trees and splitting the branch difference. Semin Cell Dev Biol. 2008;19(6):485–493. doi:10.1016/j.semcdb.2008.08.005

47. Miller KE, Suter DM. An Integrated Cytoskeletal Model of Neurite Outgrowth. Front Cell Neurosci. 2018;12. doi:10.3389/fncel.2018.00447

48. Jakobs MAH, Franze K, Zemel A. Mechanical Regulation of Neurite Polarization and Growth: A Computational Study. Biophys J. 2020;118(8):1914–1920. doi:10.1016/j.bpj.2020.02.031

49. Fan A, Tofangchi A, Kandel M, Popescu G, Saif T. Coupled circumferential and axial tension driven by actin and myosin influences in vivo axon diameter. Sci Rep. 2017;7(1):14188. doi:10.1038/s41598-017-13830-1

50. Sengupta B, Faisal AA, Laughlin SB, Niven JE. The effect of cell size and channel density on neuronal information encoding and energy efficiency. J Cereb Blood Flow Metab. 2013;33(9):1465–1473. doi:10.1038/jcbfm.2013.103

51. Kolodkin AL, Tessier-Lavigne M. Mechanisms and Molecules of Neuronal Wiring: A Primer. Cold Spring Harb Perspect Biol. 2011;3(6):a001727. doi:10.1101/cshperspect.a001727

52. Chowdhury D, Watters K, Biederer T. Synaptic recognition molecules in development and disease. Curr Top Dev Biol. 2021;142:319–370. doi:10.1016/bs.ctdb.2020.12.009

53. Südhof TC. The cell biology of synapse formation. J Cell Biol. 2021;220(7):e202103052. doi:10.1083/jcb.202103052

54. Cadart C, Heald R. Scaling of biosynthesis and metabolism with cell size. Mol Biol Cell. 2022;33(9):pe5. doi:10.1091/mbc.E21-12-0627

55. Schmoller KM, Skotheim JM. The Biosynthetic Basis of Cell Size Control. Trends Cell Biol. 2015;25(12):793–802. doi:10.1016/j.tcb.2015.10.006

56. Harrison RG. Some Unexpected Results of the Heteroplastic Transplantation of Limbs. Proc Natl Acad Sci. 1924;10(2):69–74. doi:10.1073/pnas.10.2.69

57. Penzo-Méndez AI, Stanger BZ. Organ-Size Regulation in Mammals. Cold Spring Harb Perspect Biol. 2015;7(9):a019240. doi:10.1101/cshperspect.a019240

58. Thorne RG, Nicholson C. In vivo diffusion analysis with quantum dots and dextrans predicts the width of brain extracellular space. Proc Natl Acad Sci U S A. 2006;103(14):5567–5572. doi:10.1073/pnas.0509425103

59. Kamali-Zare P, Nicholson C. Brain Extracellular Space: Geometry, Matrix and Physiological Importance. Basic Clin Neurosci. 2013;4(4):282–286.

60. Nicholson C, Hrabětová S. Brain Extracellular Space: The Final Frontier of Neuroscience. Biophys J. 2017;113(10):2133–2142. doi:10.1016/j.bpj.2017.06.052

61. Faber DS, Pereda AE. BRAIN AND NERVOUS SYSTEM | Physiology of the Mauthner Cell: Function. In: Farrell AP, ed. Encyclopedia of Fish Physiology. Academic Press; 2011:73–79. doi:10.1016/B978-0-12-374553-8.00280-X

62. Guzowski JF, Timlin JA, Roysam B, McNaughton BL, Worley PF, Barnes CA. Mapping behaviorally relevant neural circuits with immediate-early gene expression. Curr Opin Neurobiol. 2005;15(5):599–606. doi:10.1016/j.conb.2005.08.018

63. Sakurai K. Rethinking c-Fos for understanding drug action in the brain. J Biochem (Tokyo). Published online December 28, 2023:mvad110. doi:10.1093/jb/mvad110

64. Kovács KJ. Invited review c-Fos as a transcription factor: a stressful (re)view from a functional map. Neurochem Int. 1998;33(4):287–297. doi:10.1016/S0197-0186(98)00023-0

65. Dulcis D, Lippi G, Stark CJ, Do LH, Berg DK, Spitzer NC. Neurotransmitter Switching Regulated by miRNAs Controls Changes in Social Preference. Neuron. 2017;95(6):1319–1333.e5. doi:10.1016/j.neuron.2017.08.023

66. Thomas GM, Huganir RL. MAPK cascade signalling and synaptic plasticity. Nat Rev Neurosci. 2004;5(3):173–183. doi:10.1038/nrn1346

67. Hussain A, Saraiva LR, Ferrero DM, et al. High-affinity olfactory receptor for the death-associated odor cadaverine. Proc Natl Acad Sci U S A. 2013;110(48):19579–19584. doi:10.1073/pnas.1318596110

68. Cancedda L, Putignano E, Impey S, Maffei L, Ratto GM, Pizzorusso T. Patterned vision causes CRE-mediated gene expression in the visual cortex through PKA and ERK. J Neurosci Off J Soc Neurosci. 2003;23(18):7012–7020. doi:10.1523/JNEUROSCI.23-18-07012.2003

69. Ji RR, Baba H, Brenner GJ, Woolf CJ. Nociceptive-specific activation of ERK in spinal neurons contributes to pain hypersensitivity. Nat Neurosci. 1999;2(12):1114–1119. doi:10.1038/16040

70. Itoh M, Yamamoto T, Nakajima Y, Hatta K. Multistepped optogenetics connects neurons and behavior. Curr Biol CB. 2014;24(24):R1155–1156. doi:10.1016/j.cub.2014.10.065

71. Dai Y, Iwata K, Fukuoka T, et al. Phosphorylation of extracellular signal-regulated kinase in primary afferent neurons by noxious stimuli and its involvement in peripheral sensitization. J Neurosci Off J Soc Neurosci. 2002;22(17):7737–7745. doi:10.1523/JNEUROSCI.22-17-07737.2002

72. Li WC. Making In Situ Whole-Cell Patch-Clamp Recordings from Xenopus laevis Tadpole Neurons. Cold Spring Harb Protoc. 2021;2021(10). doi:10.1101/pdb.prot106856

73. Denton RD, Greenwald KR, Gibbs HL. Locomotor endurance predicts differences in realized dispersal between sympatric sexual and unisexual salamanders. Funct Ecol. 2017;31(4):915–926. doi:10.1111/1365-2435.12813

74. Offner T, Daume D, Weiss L, Hassenklöver T, Manzini I. Whole-Brain Calcium Imaging in Larval Xenopus. Cold Spring Harb Protoc. Published online October 9, 2020. doi:10.1101/pdb.prot106815

75. Padovan-Merhar O, Nair GP, Biaesch AG, et al. Single Mammalian Cells Compensate for Differences in Cellular Volume and DNA Copy Number through Independent Global Transcriptional Mechanisms. Mol Cell. 2015;58(2):339–352. doi:10.1016/j.molcel.2015.03.005

76. Voichek Y, Bar-Ziv R, Barkai N. Expression homeostasis during DNA replication. Science. 2016;351(6277):1087-1090. doi:10.1126/science.aad1162

77. Lessenger AT, Swaffer MP, Skotheim JM, Feldman JL. Somatic polyploidy supports biosynthesis and tissue function by increasing transcriptional output. Published online March 27, 2024:2024.03.25.586714. doi:10.1101/2024.03.25.586714

78. Menon T, Borbora AS, Kumar R, Nair S. Dynamic optima in cell sizes during early development enable normal gastrulation in zebrafish embryos. Dev Biol. 2020;468(1):26–40. doi:10.1016/j.ydbio.2020.09.002

79. Chen ZJ. Genetic and Epigenetic Mechanisms for Gene Expression and Phenotypic Variation in Plant Polyploids. Annu Rev Plant Biol. 2007;58:377–406. doi:10.1146/annurev.arplant.58.032806.103835

80. Yu Z, Haberer G, Matthes M, et al. Impact of natural genetic variation on the transcriptome of autotetraploid Arabidopsis thaliana. Proc Natl Acad Sci. 2010;107(41):17809–17814. doi:10.1073/pnas.1000852107

81. Osborn TC, Pires JC, Birchler JA, et al. Understanding mechanisms of novel gene expression in polyploids. Trends Genet. 2003;19(3):141–147. doi:10.1016/S0168-9525(03)00015-5

82. Desmet AS, Cirillo C, Vanden Berghe P. Distinct subcellular localization of the neuronal marker HuC/D reveals hypoxia-induced damage in enteric neurons. Neurogastroenterol Motil. 2014;26(8):1131–1143. doi:10.1111/nmo.12371

83. Fox MH. A model for the computer analysis of synchronous DNA distributions obtained by flow cytometry. Cytometry. 1980;1(1):71–77. doi:10.1002/cyto.990010114

84. Nieuwkoop PD, Faber J, eds. Normal Table of Xenopus Laevis (Daudin): A Systematical and Chronological Survey of the Development from the Fertilized Egg till the End of Metamorphosis. Garland Pub; 1994.

85. Mawaribuchi S, Takahashi S, Wada M, et al. Sex chromosome differentiation and the W- and Z-specific loci in *Xenopus laevis*. Dev Biol. 2017;426(2):393–400. doi:10.1016/j.ydbio.2016.06.015

86. Affaticati P, Le Mével S, Jenett A, et al. X-FaCT: Xenopus-Fast Clearing Technique. Methods Mol Biol Clifton NJ. 2018;1865:233–241. doi:10.1007/978-1-4939-8784-9_16

87. Preibisch S, Saalfeld S, Tomancak P. Globally optimal stitching of tiled 3D microscopic image acquisitions. Bioinformatics. 2009;25(11):1463–1465. doi:10.1093/bioinformatics/btp184

88. Schindelin J, Arganda-Carreras I, Frise E, et al. Fiji: an open-source platform for biological-image analysis. Nat Methods. 2012;9(7):676–682. doi:10.1038/nmeth.2019

89. R Core Team. R: A language and environment for statistical computing. Found Stat Comput Vienna Austria. Published online 2013.

90. Stringer C, Wang T, Michaelos M, Pachitariu M. Cellpose: a generalist algorithm for cellular segmentation. Nat Methods. 2021;18(1):100–106. doi:10.1038/s41592-020-01018-x

91. Ollion J, Cochennec J, Loll F, Escudé C, Boudier T. TANGO: a generic tool for high-throughput 3D image analysis for studying nuclear organization. Bioinformatics. 2013;29(14):1840–1841. doi:10.1093/bioinformatics/btt276

92. Arganda-Carreras I, Sorzano COS, Marabini R, Carazo JM, Ortiz-de-Solorzano C, Kybic J. Consistent and elastic registration of histological sections using vector-spline regularization. In: Proceedings of the Second ECCV International Conference on Computer Vision Approaches to Medical Image Analysis. CVAMIA’06. Springer-Verlag; 2006:85-95. doi:10.1007/11889762_8

93. Andrews S. FastQC: a quality control tool for high throughput sequence data. Published online 2010.

94. Krueger F, James F, Ewels P, Afyounian E, Schuster-Boeckler B. FelixKrueger/TrimGalore: v0. 6.7-doi via zenodo. Zenodo Doi. 2021;10.

95. Patro R, Duggal G, Love MI, Irizarry RA, Kingsford C. Salmon: fast and bias-aware quantification of transcript expression using dual-phase inference. Nat Methods. 2017;14(4):417–419. doi:10.1038/nmeth.4197

96. Soneson C, Love MI, Robinson MD. Differential analyses for RNA-seq: transcript-level estimates improve gene-level inferences. F1000Research. 2015;4:1521. doi:10.12688/f1000research.7563.2

97. Love MI, Huber W, Anders S. Moderated estimation of fold change and dispersion for RNA-seq data with DESeq2. Genome Biol. 2014;15(12):550. doi:10.1186/s13059-014-0550-8

98. Lopez V, Khakhalin AS, Aizenman C. Schooling in Xenopus laevis Tadpoles as a Way to Assess Their Neural Development. Cold Spring Harb Protoc. 2021;2021(5). doi:10.1101/pdb.prot106906

