## Supplemental data for "Ploidy and neuron size impact nervous system development and function in *Xenopus*"

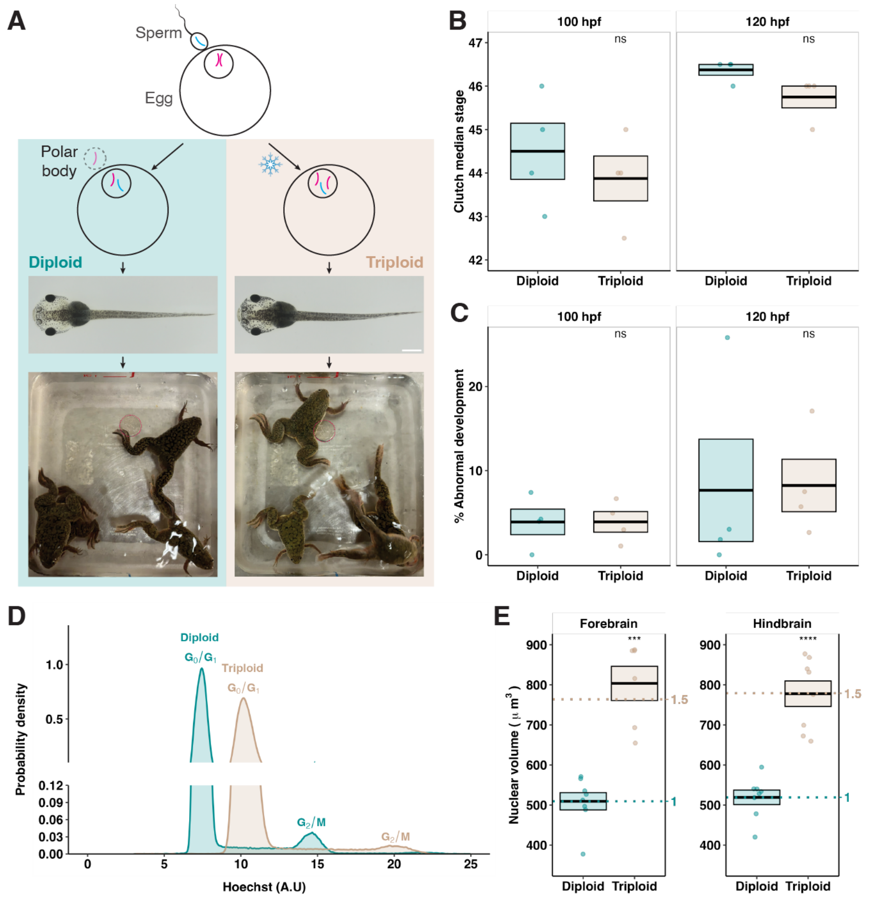


**Figure S1 Polyploid *Xenopus* as a model to study brain development**

Related to Figure 1.

**A.** Top, diagram showing *Xenopus in vitro* fertilization without or with cold shock treatment, resulting in diploid or triploid embryos, respectively[1]. Bottom, representative photos of stage 46 tadpoles (scale bar, 1 mm) and adult frogs (red dashed circle, a U.S. quarter as a size reference) of the indicated ploidy. Tadpoles and frogs shown are clutch mates. Frogs were raised by Clotilde Cadart[2]. Of note, *X. laevis* is an allotetraploid (4n=36) as a result of an ancient hybridization and genome duplication event, but has evolved into a functional diploid[3]. We refer to the natural ploidy as diploid (2n=36), and the cold-shocked, 1.5-fold ploidy as triploid (3n=54).

**B-C.** Developmental stage (B) and percentage of embryos with gross abnormalities (C) of diploid and triploid clutch mates at the indicated hours post-fertilization (hpf). Each dot represents the median of one clutch. >300 tadpoles per ploidy per time point from 4 independent clutches were examined. Abnormal-looking tadpoles were not staged. Crossbars denote mean ± SEM. ns, not significant, paired Wilcoxon signed-rank test in B and paired t test in C.

**D.** Representative density distribution of Hoechst intensity in diploid and triploid brain cells. Samples of both ploidies were obtained from the same clutch and Hoechst intensity of individual cells was analyzed by flow cytometry.

**E.** Nuclear volume of diploid and triploid neurons in the forebrain and hindbrain. Each dot represents the mean value of nuclei measured in one brain. 8 brains per ploidy across 3 independent clutches were examined. Crossbars indicate mean ± SEM and dotted lines mark 1- and 1.5-fold of diploid mean. ***, $p<0.001$; ****, $p<0.0001$, t test.

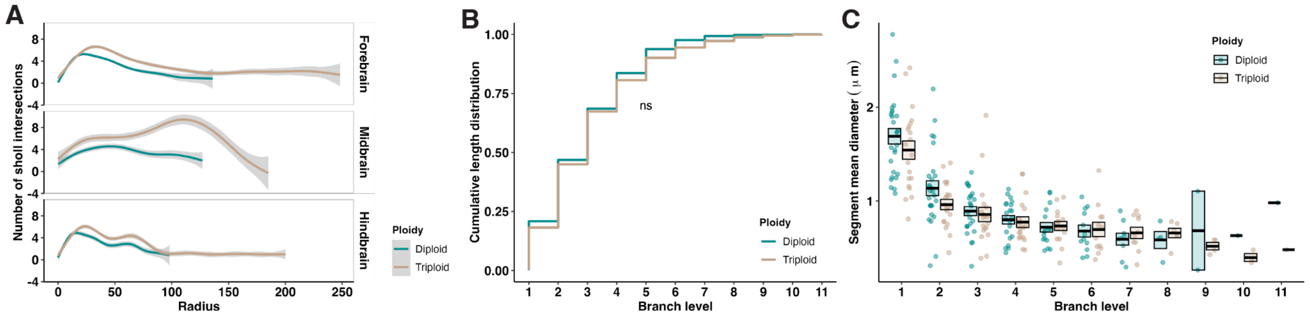


**Figure S2 Size and shape analysis of diploid and triploid neurons**

Related to Figure 1.

**A.** Sholl analyses of diploid and triploid neurons in the indicated brain regions. The same neurons were used as in Figures 2C. Data were smoothed with a generalized additive model (GAM) and were presented as mean ± 95% confidence interval.

**B.** Cumulative length distribution of neurites at different branch levels. ns, not significant, Kolmogorov-Smirnov test.

**C.** Mean neurite segment diameter at different branch levels. Same data as in Figure 1I.

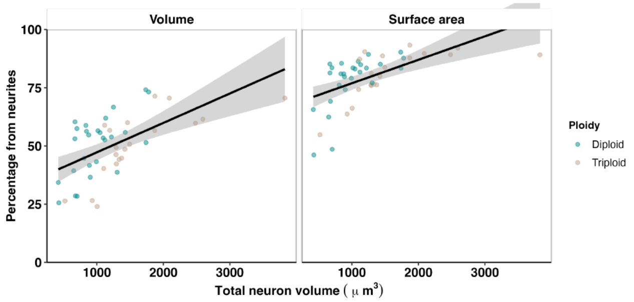


**Figure S3 Proportion of neuronal volume and surface area contributed by neurites**

Related to Figure 2.

Diploid and triploid data were combined, smoothed with a linear model, and presented as mean ± 95% confidence interval.

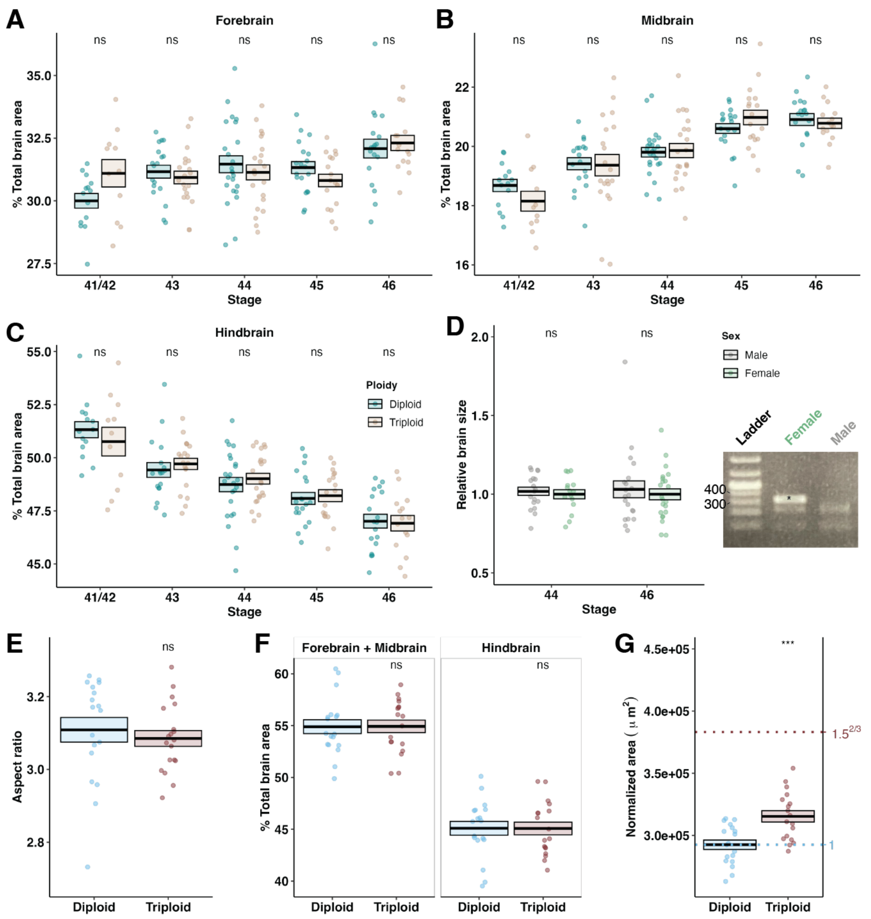


**Figure S4 Triploid brains are morphologically similar to diploid brains**

Related to Figure 3.

**A-C.** Comparison of the proportion of the forebrain (A), midbrain (B), and hindbrain (C) in diploid and triploid brains across multiple developmental stages. Each dot represents one brain. Numbers of diploid/triploid brains examined were 15/11 at stage 41/42, 18/20 at stage 43, 24/22 at stage 44, 19/18 at stage 45, and 18/15 at stage 46. Brains were from 3 independent clutches.

**D.** Relative size of female and male diploid brains. Each dot represents one brain. Numbers of male/female brains examined were 15/18 at stage 44 and 22/20 at stage 46. Brains were from 3 independent clutches. Brain size was divided by its clutch female mean to normalize against clutch variance. Insert on the bottom right shows an example DNA gel used to determine the sex of tadpoles. The presence of a W chromosome-specific amplicon (315 bp, asterisk) indicates a female[4].

**E-G.** Comparisons of the aspect ratio (E), proportion of different brain regions (F), and normalized area (G) of diploid (2n=20) and triploid (3n=30) *X. tropicalis* brains at stage 46. Each dot represents one brain. 18 brains per ploidy from 3 independent clutches were examined.

In A-G, crossbars denote mean ± SEM. *, p<0.05; **, p<0.01; ***, p<0.001; ****, p<0.0001; ns, not significant, t-test.

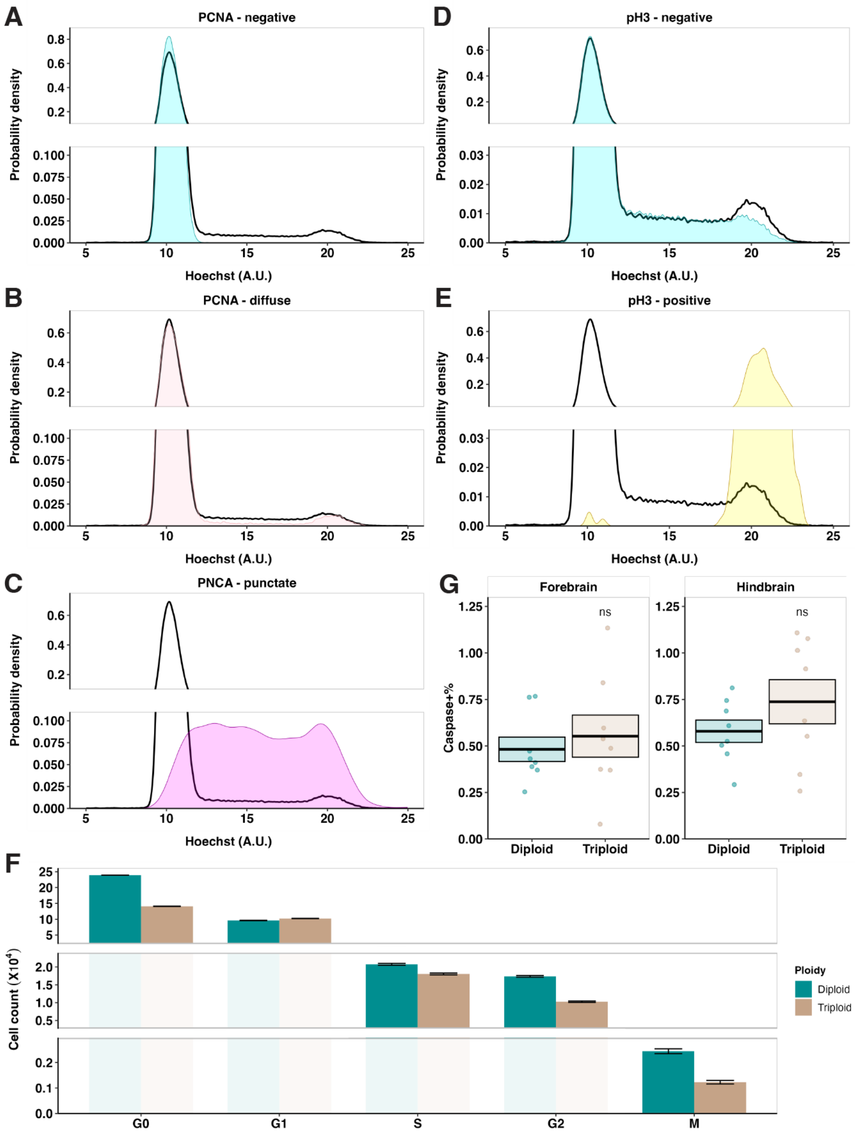


**Figure S5 Triploid brains show distinct cell birth/death dynamics**

Related to Figure 4.

**A-E.** Probability density distributions of Hoechst intensity of the three different PCNA populations (A-C) and the two different pH3 populations (D-E) in a representative flow cytometry sample of dissociated brain cells at developmental stage 46. Colored curves show the distribution of the indicated population and black curves show the distribution of the total population as a reference.

**F.** Comparison of cell counts in different cell cycle phases at developmental stage 46. Error bars mark 95% confidence interval. Same data as in Figure 4G.

**G.** Ratio of cell death (marked by the positive staining of cleaved caspase-3[5]) in the indicated brain region in diploid and triploid brains. Number of caspase-positive cells was normalized to total cell count in the same region. Each dot represents one brain. 8 brains per ploidy across 3 independent clutches were examined. Crossbars denote mean ± SEM. ns, not significant, t-test.

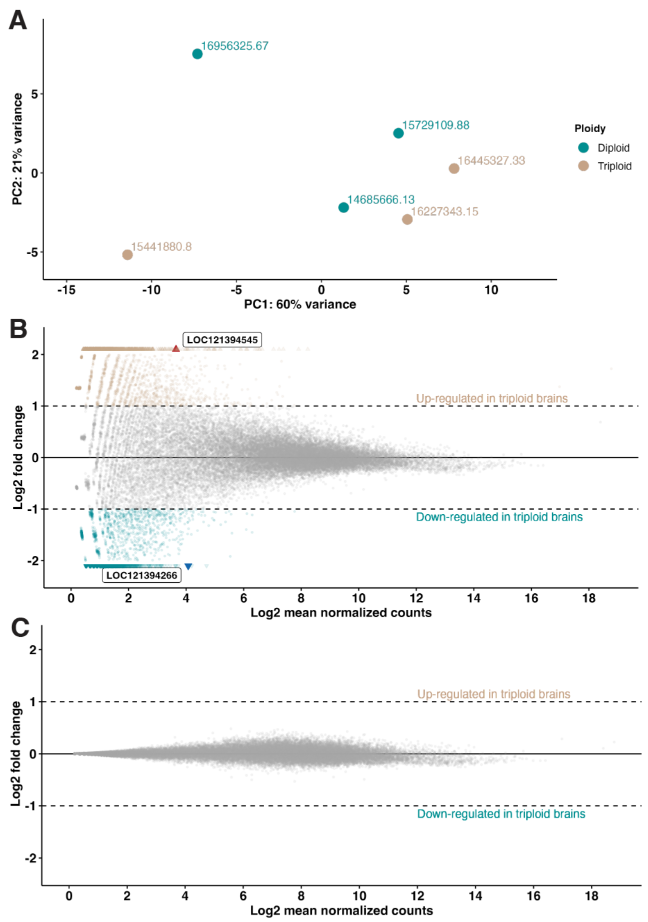


**Figure S6 Triploid and diploid brains possess similar transcriptional profiles**

Related to Figure 4.

**A.** Principal component analysis (PCA) for brain-specific RNA-seq replicates. 3 replicates per ploidy were used and each replicate was prepared from >10 live-dissected brains of clutch-controlled, anesthetized stage 44-46 tadpoles. Total normalized count is shown for each replicate.

**B.** Mean-Average (MA) plot showing log2 fold changes (triploid/diploid) of gene expression in tadpole brains (normalized by distribution). Points that fall outside of the y-axis limits are plotted as triangles. The two annotated points are the only two with adjusted $p<0.1$ and are both uncharacterized genes. Red triangle, LOC121394545, is predicted to encode E3 ubiquitin-protein ligase DCST1-like, and blue triangle, LOC121394266, is predicted to encode general transcription factor II-I repeat domain-containing protein 2-like (Xenbase *Xenopus laevis* J strain 10.1).

**C.** MA plot with log2 fold changes in B moderated with normal shrinkage to remove noise[6]. No point falls outside of the y-axis limits. No point has adjusted $p<0.1$.

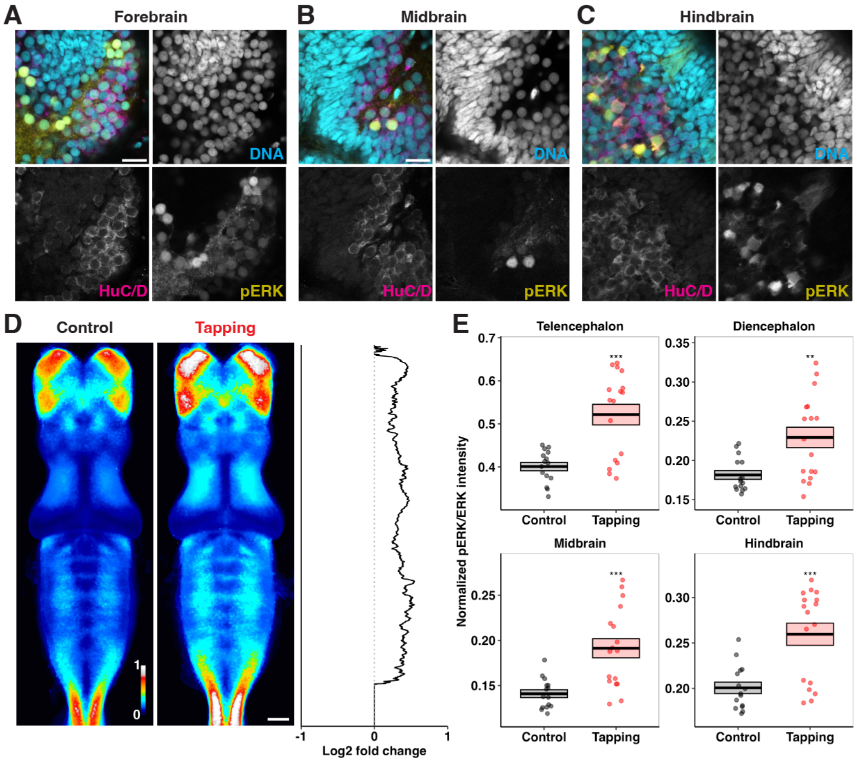


**Figure S7 pERK expression is neuronal and reflects neural activity**

Related to Figure 5.

**A-C.** Representative micrographs of pERK and HuC/D co-staining in the forebrain (A), midbrain (B), and hindbrain (C) of stage 46 tadpoles. Scale bar, 20 μm.

**D.** Heatmap showing normalized, averaged pERK/ERK intensity of 15 control brains and 17 brains of tadpoles after 15 min of dish tapping. Stage 46 diploid tadpoles from 3 independent clutches were used. Log2 fold changes (tapping/control) of pERK/ERK intensity along Y axis are plotted to the right. Scale bar, 100 μm.

**E.** Comparison of pERK/ERK intensity in the indicated brain regions. Each dot represents one brain. Crossbars denote mean ±SEM. **, $p<0.01$; ***, $p<0.001$, t test.

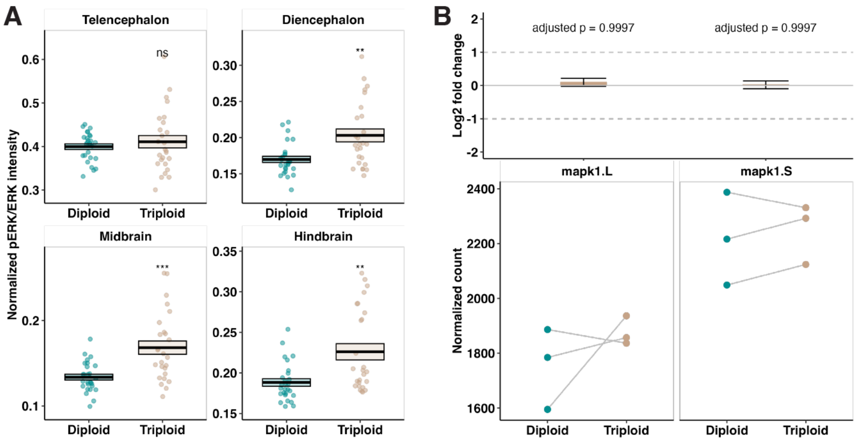


**Figure S8 Triploid brains show increased pERK/ERK level but unaltered ERK expression**

Related to Figure 5.

**A.** Comparison of pERK/ERK intensity in the indicated brain regions. Same data as in Figure 5A-B. Each dot represents one brain. Crossbars denote mean ±SEM. **, $p<0.01$; ***, $p<0.001$; ns, not significant, t test.

**B.** Normalized counts and log2 fold changes (triploid/diploid) of the two ERK genes in *X. laevis* (L and S homeologs). Grey lines connect samples from the same clutch. This is from the same brain-specific RNA-seq as in Figures S6.

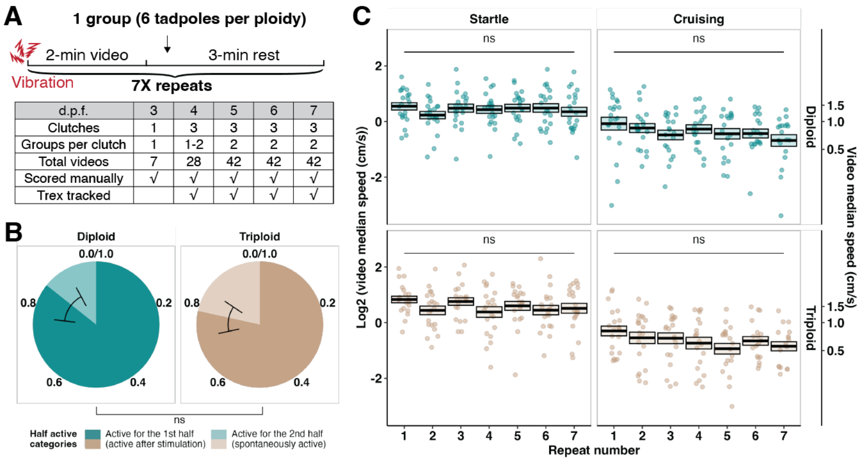


**Figure S9 Assessing stimulated swimming, spontaneous swimming, and learning behavior**

Related to Figure 6.

**A.** Diagram showing the time scheme and number of replicates used for the swimming assay.

**B.** Break-down of the “half active” category in Figure 7B. Error bars mark 95% confidence interval. ns, not significant, Fisher’s exact test.

**C.** Swimming speeds of tadpoles as they experienced repeated stimulation. Each dot represents the geometric mean of all TRex-tracked speeds in one video. Data from 4-7 dpf were pooled for this analysis. Crossbars denote mean ± SEM. ns, not significant, ANOVA test.

**REFERENCES**


1. Gibeaux R, Heald R. Generation of Xenopus Haploid, Triploid, and Hybrid Embryos. *Methods Mol Biol Clifton NJ*. 2019;1920:303-315. doi:10.1007/978-1-4939-9009-2_18

2. Cadart C, Bartz J, Oaks G, Liu MZ, Heald R. Polyploidy in *Xenopus* lowers metabolic rate by decreasing total cell surface area. *Curr Biol*. 2023;33(9):1744-1752.e7. doi:10.1016/j.cub.2023.03.071

3. Session AM, Uno Y, Kwon T, et al. Genome evolution in the allotetraploid frog Xenopus laevis. *Nature*. 2016;538(7625):336-343. doi:10.1038/nature19840

4. Mawaribuchi S, Takahashi S, Wada M, et al. Sex chromosome differentiation and the W- and Z-specific loci in *Xenopus laevis*. *Dev Biol*. 2017;426(2):393-400. doi:10.1016/j.ydbio.2016.06.015

5. Crowley LC, Waterhouse NJ. Detecting Cleaved Caspase-3 in Apoptotic Cells by Flow Cytometry. *Cold Spring Harb Protoc*. 2016;2016(11). doi:10.1101/pdb.prot087312

6. Love MI, Huber W, Anders S. Moderated estimation of fold change and dispersion for RNA-seq data with DESeq2. *Genome Biol*. 2014;15(12):550. doi:10.1186/s13059-014-0550-8
